# Minimising severity of dengue serotype 1 infection by transmissible interfering particles

**DOI:** 10.1101/2020.04.21.052936

**Authors:** A. Shausan, J. Aaskov, C. Drovandi, K. Mengersen

## Abstract

Transmissible interfering dengue particles (DENV–TIPs) are engineered dengue virus mutants which are defective and can replicate only with the help of dengue wild–type virus (DENV). *In vitro* studies have found that when DENV–TIPs and DENV coinfect a cell, they compete for viral genomes and cell proteins for replication and packaging, and DENV–TIPs outperform DENV in this process. Thus, it is hypothesised that DENV–TIPs may be used as a novel therapeutic agent. However, the effectiveness of DENV–TIPs as an antiviral agent is yet to be explored at an epidemiological scale. We present a mathematical model for the replication of DENV and DENV–TIPs as they interact with human host cells, accounting for the effectiveness of DENV–TIPs in blocking DENV from coinfected cells. We fit the model to sequentially measured plasma viral titre data from primary and secondary dengue serotype 1 infected patients in Vietnam. We show that variation in initial DENV load is sufficient to recreate the observed variation between patients. Parameter estimates, differing in primary and secondary infections, do not confirm a significant difference between these two types of infection. We use our model to investigate the potential impact of DENV–TIPs as an antiviral agent. We conclude that, when the effectiveness of DENV–TIPs in inhibiting DENV from coinfected cells is at least 80%, a dose as high as 10^12^ copies per millilitre of blood is required to reduce duration of infection and peak DENV serotype 1 infection level at any time point of infection. This work provides a quantitative understanding of the relationship between DENV–TIPs levels and their efficiency in clearing dengue viral infection. It will guide future development of mechanistic models of how DENV–TIPs might contribute as an antiviral agent in limiting natural dengue infection.

**Author summary:** Inhibition of dengue wild–type virus (DENV) by transmissible interfering dengue particles (DENV–TIPs) is seen in some *in vitro* studies, and it is hypothesised that DENV–TIPs may be used as a therapeutic agent. However, the efficiency of DENV–TIPs in limiting DENV infection in patients is yet to be explored at an epidemiological scale. Using data collected from dengue serotype 1 infected patients, we model how DENV replicates in an infected patient and how effective DENV–TIPs are in controlling that replication. Our results are of use in the evaluation of DENV–TIPs as a potential antiviral agent.

## Introduction

Dengue is a mosquito–borne viral disease in humans, with an estimated prevalence of 390 million per year, of which roughly 25% show clinical symptoms [1]. Approximately 3.5 billion people, in 128 countries, are at risk of dengue infection [2, 3]. The transmission of dengue virus involves the *Aedes aegypti* mosquitoes as the primary vectors, and humans as the main source and victims of infection [4]. Human infection is acute and self–limiting and may range from being mild to one of the severe forms, such as dengue hemorrhagic fever or dengue shock syndrome, which may be life–threatening.

There are four distinct serotypes of dengue wild–type virus (DENV1–4). In humans, recovery from infection by one serotype provides lifelong immunity against that particular serotype. However, cross–immunity to other serotypes after recovery is only partial and temporary. Successive secondary infections by a heterologous serotype increase the risk of developing severe dengue [5, 6]. The cause of this is envisaged as the insufficiency of the antibodies generated in the primary infection to neutralise the virus from secondary infection, but still attach to virus particles. The resulting partially neutralised immune complexes are thought to increase viral replication process. This hypothesis is known as the antibody–dependent enhancement (ADE) [7–9].

Current dengue control programs emphasise either killing the *Aedes* vector or reducing the breeding of larvae by removing breeding sites and/or killing the larvae. There is little evidence that these measures effectively reduce the burden of dengue on a global scale due to their huge expense and the negative impact on human health, for example by the application of insecticides such as Dichlorodiphenyltrichloroethane [10]. An alternative experimental control strategy is to minimise the lifespan of adult *Aedes aegypti* mosquitoes by infecting them with the bacteria *Wolbachia* [11]. However, application of this process in the field is not straightforward [12]. Vaccination of humans is another control strategy. Nevertheless, the only licensed dengue vaccine, *Dengvaxia*, requires three doses of vaccine over 12 months, and cannot be used for children under the age of 9 years or in areas where pre–existing anti–DENV antibody prevalence is less than 70 per cent [13, 14]. Two other vaccines under development (N.I.H., Takeda) may not require multiple injections to reach similar or higher levels of protection but their success is untested [14]. Thus, alternative methods to combat the challenges caused by vector control and vaccination programs are required.

It is shown that increases in a patient’s plasma DENV level is associated with a higher probability of human to *Aedes aegypti* transmission of DENV, and the likelihood of successful transmission coincides with a patient’s febrile period and the dynamics of DENV level [15]. Thus, any reduction in a patient’s plasma DENV and/or duration of infection may limit the chance of human to mosquito transmission, which may then disrupt the human–mosquito–human transmission cycle of DENV.

A proposed strategy to decrease virus level in humans is the use of *transmissible interfering* DENV *particles* (DENV–TIPs) as a therapeutic agent [16]. DENV–TIPs are engineered DENV mutants which are non–infectious and lacking part of the essential genome for replication. TIPs utilise the concept of naturally occurring *defective interfering particles* (DIPs), which appear in many RNA virus infections, such as influenza, HIV/AIDS and dengue [17]. DIPs are unable to replicate unless the cell which they infect also contains wild–type virus (WT–virus), which contribute genes which are missing or defective in DIPs. Within a coinfected cell, DIPs and WT–virus compete for viral proteins in order to replicate and package and ultimately, DIPs outperform WT–virus, producing DIPs mostly from the cell [18, 19]. Inhibition of WT–virus replication by DIPs is first identified more than 60 years ago with influenza virus [20]. Since then, many *in vitro* studies have investigated the mechanism which DIPs inhibit WT–virus replication [17, 21–25], and the potential for utilising DIPs as an anti–viral agent [16, 26, 27].

A recent study has found that infecting *Aedes aegypti* mosquitoes with blood sera from DENV infected patients produced DIPs in mosquitoes which are identical to those observed in patients, suggesting that DIPs may be transmitted between humans and mosquitoes [28]. This study has also established that engineered DIPs (DENV–TIPs), reduced the yield of DENV following coinfections of susceptible cells *in vitro*. These findings suggest the possibility of DENV–TIPs as an anti–viral agent for dengue patients and their transmissibility to mosquitoes. The effectiveness of DENV–TIPs as a therapeutic agent at the epidemiological scale is yet to be explored.

Given that the plasma virus level and patient’s febrile period are indicators of successful DENV transmission to mosquitoes [15], and that DENV–TIPs have the potential to act as an anti–viral agent [28], it is important to investigate the dynamics of DENV–TIPs and DENV as they interact with human hosts, and assess the effectiveness of DENV-TIPS in shaping a patient’s virus profile. Mathematical modelling of the interaction of DENV and immune response, validated against patient data, have proved a useful tool for gaining an understanding of the role which immunity plays in shaping patients’ DENV dynamics [29–32]. However, these models do not describe the dynamics of DENV–TIPs. There are few models which describe the dynamics of DIPs and DENV [33–35], and only one [35] amongst these targets within human host dynamics, but this is not validated against patient data. Others have aimed to study the kinetics occurring in *in vitro* experiments [33, 34]. None of these previous models involving DIPs have explicitly accounted for effectiveness of DIPs in limiting DENV infection.

We develop a mathematical model of dengue for the dynamics of DENV–TIPs and DENV as they interact with human host cells, explicitly accounting for effectiveness of DENV–TIPs in blocking DENV production from coinfected cells. We validate the model using sequentially measured viral titre data from DENV1 infected patients, who showed clinically apparent symptoms. The resulting parameter estimates guide the determination of factors related to the variability observed in infection dynamics between patients, but do not suggest a difference between primary and secondary DENV1 infections. By using the estimated parameters we perform two simulation studies to examine the potential impact of DENV–TIPs therapy. Our analyses allow us to hypothesise a minimal DENV–TIPs dose required and an indication of how effective DENV–TIPs must be in order to limit disease severity in DENV1 infected patients.

## Materials and methods

### The data

We use patient–level viral load data to validate the mathematical model. The data are obtained from a previous experimental study on DENV transmission from humans to mosquitoes [15]. The study in [15] concerns 208 adult dengue patients who visited the Hospital for Tropical Diseases in Ho Chi Minh City, Vietnam. Patients were enrolled within 72 hours of fever and experimentally exposed to field–derived *Aedes aegypti* mosquitoes on 2 randomly chosen days during their first 4 days in the study. Virus RNA in plasma was quantified by RT–PCR assay and each patient had once daily virus measurement recorded for a maximum of 5 days. Measurements were recorded as copies per millilitre (copies/ml) of plasma and patients were categorised according to their infecting serotype (DENV1–4) and serology (primary, secondary). Serology of some patients was not identified and thus marked as *indeterminate*. See [15] for details on classifications of serotype and serology. The RT–PCR assay used had a limit of detection (LOD) for each serotype, with 357 copies/ml as the LOD for both DENV1 and DENV3, 72 copies/ml for DENV2 and 720 copies/ml for DENV4.

We use data on DENV1 primary (*n* = 25) and secondary (*n* = 42) patients in our analyses (Fig 1). There were 11 DENV1 cases whose serotype was *indeterminate*, so we exclude these cases for model fitting.

**Fig 1.**
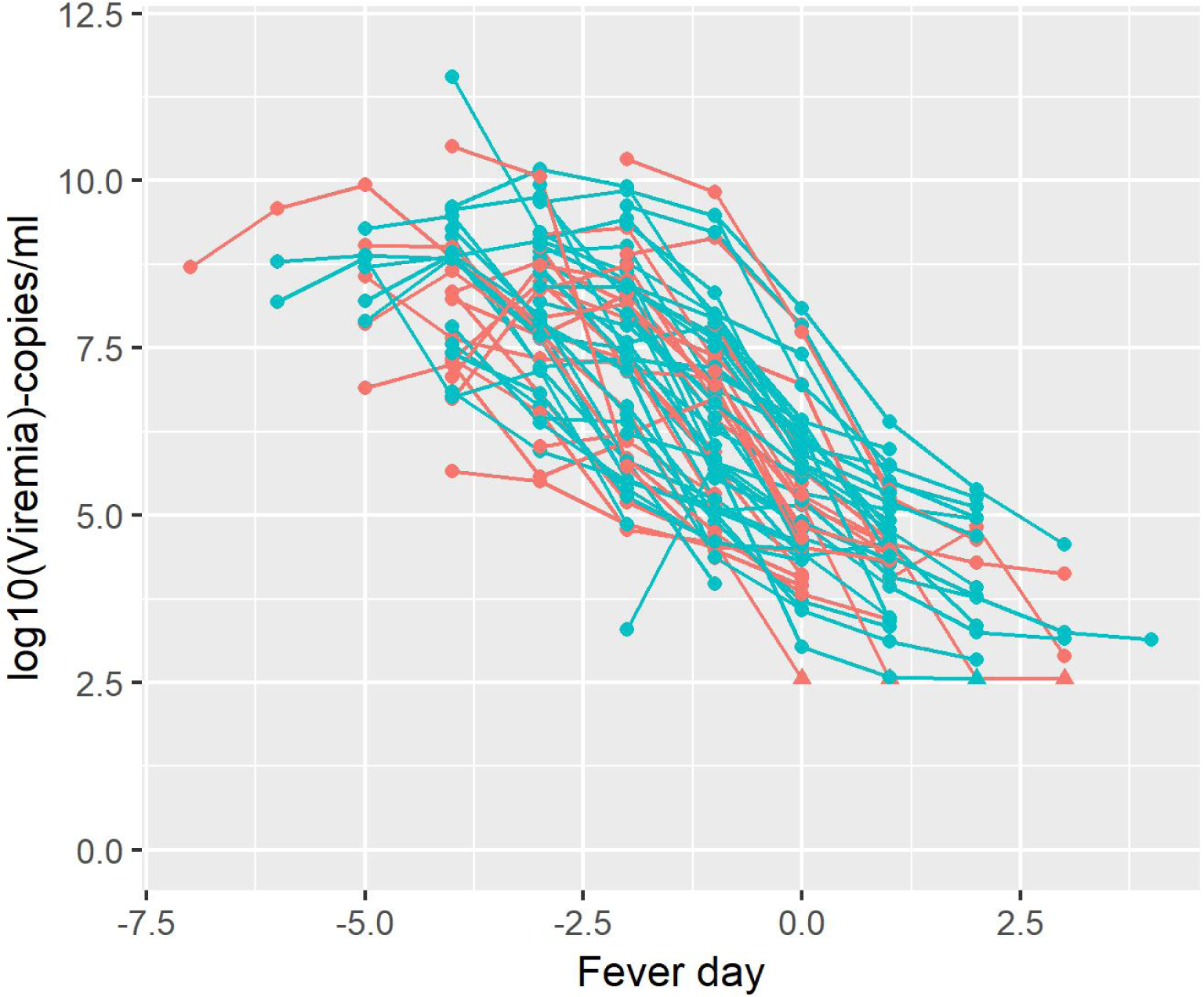
Plot of DENV1 profiles by fever day. Colours represent primary (red) and secondary (blue) profiles. Circles and triangles resemble measurements above and below LOD, respectively. Fever day 0 corresponds to the day of defervescence.

### Mathematical model definition

Given that there are no experimental results determining how effective DENV–TIPs are in shaping DENV dynamics of patients, we propose a novel within–host mathematical model for the replication of DENV–TIPs and DENV as they interact with human host cells, accounting for effectiveness of DENV–TIPs in blocking DENV production from coinfected cells. The model is constructed using the within–host DIPs and WT–virus replication process suggested in [18] and [36].

The model is a system of ordinary differential equations (ODE) with six state variables: the population sizes of uninfected susceptible cells, *S*(*t*); cells infected with only DENV, *I*_*V*_(*t*), only DENV–TIPs, *I*_*T*_(*t*), both DENV and DENV–TIP, *I*_*B*_(*t*) (hereafter referred as *coinfected cells*); free DENV, *V* (*t*); and free TIPs, *T* (*t*), at time *t*.

The model is described by the following equations:

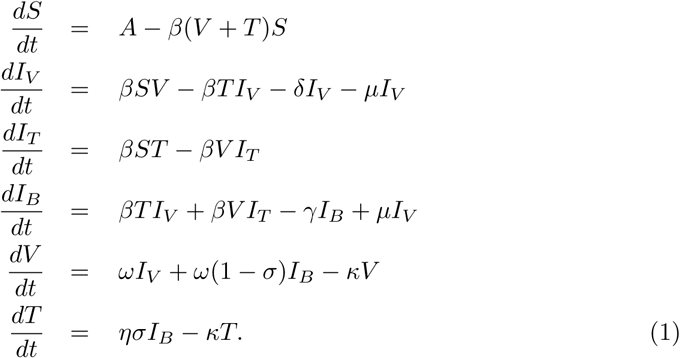

An uninfected Susceptible cell is created at a constant rate, *A*, and can become infected by a DENV or a DENV–TIP particle at rate *β*, via a mass action process, converting it to an *I*_*V*_ or *I*_*T*_ cell, respectively. An *I*_*B*_ cell is created when a DENV-TIP particle infects an *I*_*V*_ cell or a DENV particle infects an *I*_*T*_ cell at rate *β*, via a mass action process, or a DENV particle mutates within an *I*_*V*_ cell to produce a DENV–TIP at rate *µ*. Parameters *δ* and *γ* represent death rates of an *I*_*V*_ and *I*_*B*_ cell, respectively. A DENV or a DENV–TIP particle dies at rate *κ*. An *I*_*V*_ type cell can produce only DENV particles and the rate of this production is *ω*. An *I*_*T*_ type cell does not produce any particles. It is assumed that there is a one-to-one competition between DENV–TIPs and DENV particles within an *I*_*B*_ cell for viral proteins, and this competition may inhibit DENV production fully or partially from the cell. Let parameter *σ* ∈ [0, 1] be an indication of effectiveness of DENV–TIPs in blocking DENV production in this competition. If *σ* = 1, the presence of at least one DENV–TIP fully inhibits DENV production from an *I*_*B*_ cell and only DENV–TIPs are produced at rate *η*. If *σ* = 0, none of the DENV–TIPs can inhibit virus production and the cell produces only DENV at rate *ω*. If DENV–TIPs are partially effective, in which case *σ* ∈ (0, 1), the *I*_*B*_ cell produces both DENV–TIPs and DENV. However, the rate of DENV production from the cell is reduced by a factor of *ω*(1 − *σ*).

Our ODE model 1 has some similarities and differences from the within–host models constructed in [34] and [35], which describe the dynamics of WT–virus and DIPs as they interact with human host cells. Like our model, the model studied in [34] is a system of ODE, but considers the ages of cells infected with only WT–virus and coinfected cells, by incorporating subclasses corresponding to stages through which these cells pass until cell burst time is reached. The model proposed in [35] is a system of partial differential equations considering a spatial component for abundance of viral and cell types, adding a non-–mass action process of mixing viral and cell components, and incorporating age of cells infected with only WT—type virus. Neglecting age of cells infected with DENV and DENV—TIPs in our model is an obvious simplification, but in the absence of detailed data on cell concentration and ages of cells to fit the model to, a more complex description cannot be parametrised. We do not distinguish between DENV–TIPs and DIPs in our ODE, as DENV–TIPs are engineered from naturally arising DIPs [28], and we assume the kinetic of DENV–TIPs within human hosts will be the same as that of DIPs.

Model (1) differs from the within–host dengue models analysed in [31, 32, 37], in which the authors assumed that clearance of infected cells or free virus are mediated by antibodies. Model (1) also differs from the models studied in [29, 30], which examined the roles of both innate and adaptive immunity play in clearing infected cells. None of these models accommodated the dynamics of DENV–TIPs. As our focus is primarily on examining the role which DENV–TIPs play in shaping DENV dynamic within patients, we model neither innate nor adaptive immunity.

### Parameter estimation

Model (1) contains 9 parameters (*A, β, δ, µ, γ, ω, κ, σ*) and 6 initial conditions 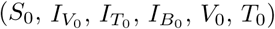. Table 1 provides the meaning of these parameters and initial conditions, specifies whether they are estimated or assigned, and shows nominal values used for parameters with reference where appropriate.

**Table 1.**
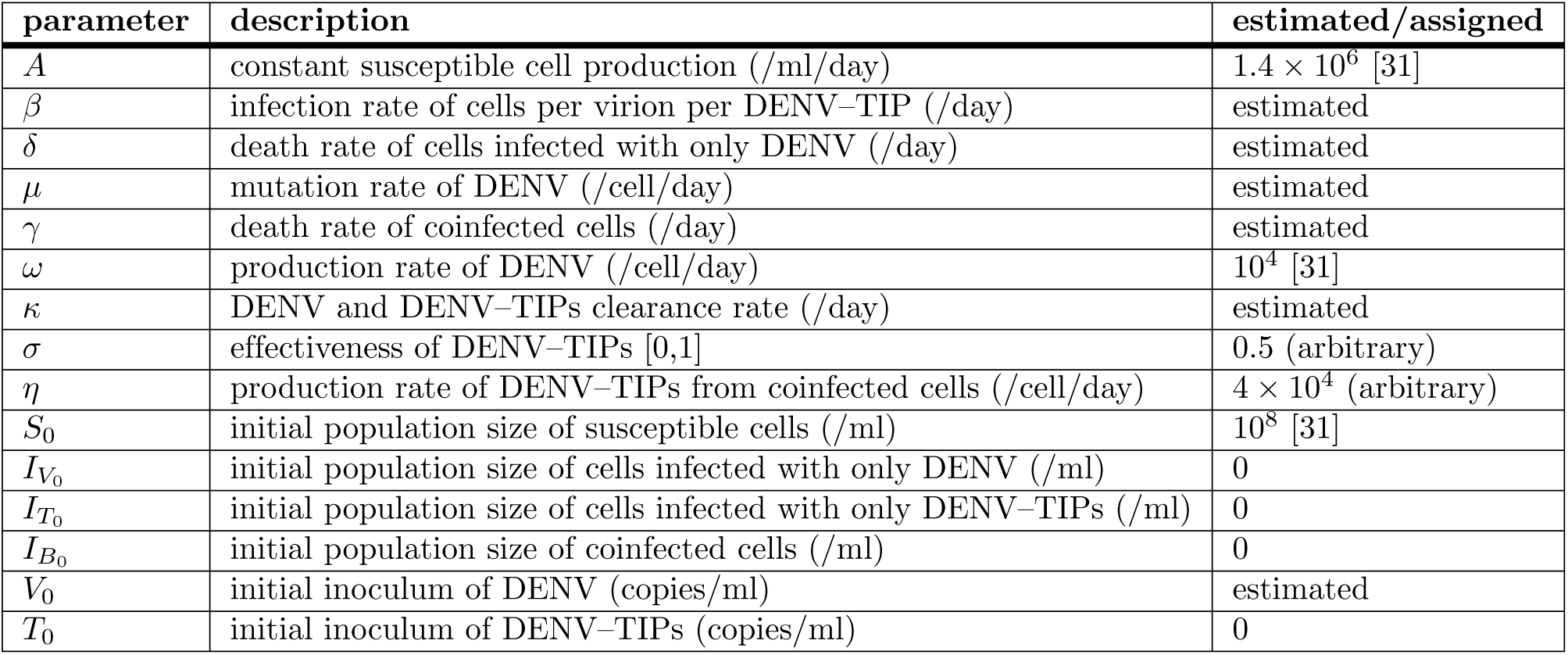
The parameters of the model and values if assigned.

| parameter | description | estimated/assigned |
| --- | --- | --- |
| $A$ | constant susceptible cell production (/ml/day) | $1.4 \times 10^6$ [31] |
| $\beta$ | infection rate of cells per virion per DENV-TIP (/day) | estimated |
| $\delta$ | death rate of cells infected with only DENV (/day) | estimated |
| $\mu$ | mutation rate of DENV (/cell/day) | estimated |
| $\gamma$ | death rate of coinfecting cells (/day) | estimated |
| $\omega$ | production rate of DENV (/cell/day) | $10^4$ [31] |
| $\kappa$ | DENV and DENV-TIPs clearance rate (/day) | estimated |
| $\sigma$ | effectiveness of DENV-TIPs [0,1] | 0.5 (arbitrary) |
| $\eta$ | production rate of DENV-TIPs from coinfecting cells (/cell/day) | $4 \times 10^4$ (arbitrary) |
| $S_0$ | initial population size of susceptible cells (/ml) | $10^8$ [31] |
| $I_{V_0}$ | initial population size of cells infected with only DENV (/ml) | 0 |
| $I_{T_0}$ | initial population size of cells infected with only DENV-TIPs (/ml) | 0 |
| $I_{B_0}$ | initial population size of coinfecting cells (/ml) | 0 |
| $V_0$ | initial inoculum of DENV (copies/ml) | estimated |
| $T_0$ | initial inoculum of DENV-TIPs (copies/ml) | 0 |

The reason for assigning some parameters nominal values is because, in the absence of data on concentration of DENV–TIPs and the four types of cells considered in the model, not all parameters are identifiable. This is seen by performing an identifiablity analysis, similar to procedures outlined in [30, 32]. Substituting *Ŝ* = *S/A, Î*_*V*_ = *I*_*V*_ */A, Î*_*T*_ = *I*_*T*_ */A, Î*_*B*_ = *I*_*B*_*/A*, 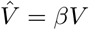 and 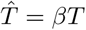, in the ODE (1) gives:

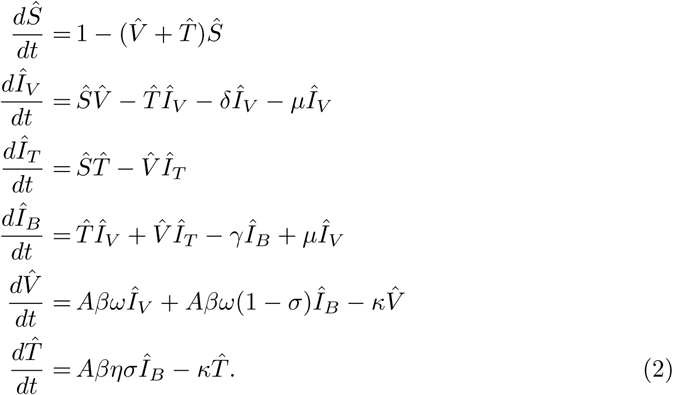

These equations reveal that parameters appearing in the sets *Aβω*(1 − *σ*) and *Aβησ* cannot be uniquely identified. As measurements of DENV level for each patient are available, *β* and initial DENV load (*V*_0_) can be estimated independently. Therefore, we fix parameters *A, ω, σ* and *η* at nominal values and estimate *β, δ, µ, γ, κ* and *V*_0_. Estimated parameters are fitted either common to all patients, specific to primary and secondary groups, or specific to each patient.

We assign values for the rates of susceptible cell production (*A*), DENV production (*ω*), and initial population size of susceptible cells (*S*_0_) based on [31]. We assume that patients’ blood does not contain any infected cells or free DENV–TIPs, at the time they become infected with free DENV. Hence, we set the initial population sizes of cells infected with only DENV 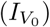, only DENV–TIPs 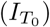, both DENV and DENV–TIPs 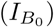, and initial inoculum of DENV–TIPs (*T*_0_) to 0. Due to a lack of knowledge of both the effectiveness of DENV–TIPs in blocking DENV from coinfected cells (*σ*) and rate of production of DENV–TIPs from coinfected cells (*η*), we assign arbitrary values for these two parameters. We discuss the sensitivity of the assigned parameters to DENV output, from the ODE, in the *Results* section.

The only initial condition we fit in our statistical analysis is the initial amount of free DENV (*V*_0_), which we estimate as patient–specific. This is biologically intuitive, as each patient may receive a different amount of DENV by infected mosquitoes when they take blood meals from people and mosquitoes may be interrupted during feeding.

Parameters *β, δ, µ, γ, κ* are fitted either common to all patients (table 2, model *a*) or specific to primary and secondary groups (table 2, model *b*). Our goal in fitting these two model variants is to determine which of these hypotheses best account for the observed variation in viral load patterns in primary and secondary DENV1 infected patients (Fig 1). Previous dengue studies allowed either the virus replication rate (*β*) [29–32] or both *β* and the virus clearance rate (*κ*) [32] to vary by primary and secondary infections, to test the ADE process. However, our aim in using model *b* is not specifically to investigate the ADE process, but to test any potential biological differences between primary and secondary infections which may be associated with DENV–TIPs.

**Table 2.** The parameters of the model and values if assigned. Common parameters are shared across all patients regardless of their group (primary or secondary). Also shown are the WAIC value for each model and its standard error.

| model type | common parameters | group-specific parameters | patient-specific parameters | WAIC (SE) |
| --- | --- | --- | --- | --- |
| *a* | $\beta, \delta, \mu, \gamma, \kappa$ | – | $V_0$ | 11600 (225.5) |
| *b* | – | $\beta, \delta, \mu, \gamma, \kappa$ | $V_0$ | 11608 (225.7) |

The DENV1 data are available as a function of time since the onset of symptoms. However, we intend to model DENV dynamic from the start of infection. To facilitate this, it is required to estimate the incubation period (IP) or equivalently, the initial time of infection. IP is defined as the time between DENV inoculation and the start of symptoms. We expect the IP to correlate with the initial amount of free DENV (*V*_0_), which we estimate. Thus, we do not aim to estimate the IP. Instead, following [30], we use the estimate derived in [38] which used Bayesian time–to–event models to estimate IP using observations from 35 dengue studies. This estimate is 5.9 days with 95% confidence interval lying between 3 and 10 days. We add 6 days, which is the estimated IP rounded to nearest whole number, prior to the first measurement of each patient. This amounts to starting infection randomly, for each patient.

We use a Bayesian approach for parameter estimation. Based on the dynamics of virus profiles given in Fig 1 and following [30, 32], we hypothesise that, for each patient, the viremia measurements follow a log–normal distribution with mean given by the logarithm of virus output from the ODE (1). The likelihood for the *j*–th patient is:

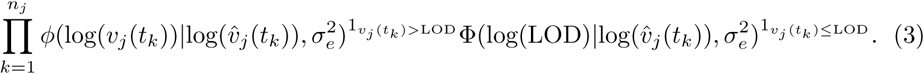

In the likelihood, *n*_*j*_ denotes the number of virus measurements for the *j*–th patient, and *v*_*j*_(*t*_*k*_) and 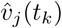 are the observed and modelled virus measurements, respectively, for the *j*–th patient at time *t*_*k*_. We use a Gaussian probability density function, *ϕ*, to calculate the probability of observing a virus measurement only if it lies above the LOD. For measurements below the LOD, all we know is that they are at or below the LOD. Thus, we use the normal cumulative distribution function, Φ, to calculate the probability of observing a data point at or below the LOD. The parameter quantifying the standard deviation of the measurement error is *σ*_*e*_, which we estimate here. The full log–likelihood is the sum of the log–likelihoods over all patients.

In both models *a* and *b*, prior distributions for *β, δ, µ, γ, κ* are assumed to be uniformly distributed with ranges chosen to include values which are previously established and biologically plausible. The range set for *δ, κ, γ*, is from 1 to 10, which includes the estimated value of 3.30 with credible interval of (3.07, 3.48) for *δ* and the assigned value of 3.5 for *κ* in [31] for their virus neutralisation model. We expect the value of *γ* to be close to but less than *δ* and thus seek an estimate of it within the same range as *δ*. The range for *β* is from 2 × 10^−11^ to 2 × 10^−10^, which includes the values of 3.83 × 10^−11^ and 5 × 10^−11^ assigned for primary and secondary infections, respectively, in [31] for their virus neutralisation model. For *µ*, the range set is from 5 × 10^−5^ to 1 × 10^−3^, which includes the value of 1 × 10^−4^ used in [35]. The prior distribution for *V*_0_ is assumed vague relative to the data and anticipated values, so is set to be normally distributed with mean 0 and standard deviation of 100. Similarly, the prior distribution for *σ*_*e*_ is assumed normally distributed with mean 1 and standard deviation of 2.

We use a Markov Chain Monte Carlo (MCMC) algorithm implemented in the probabilistic programing language *Stan* for parameter estimation, Specifically, the no–U–turn sampler (NUTS), which is a variant of the Hamiltonian Monte Carlo (HMC) algorithm, is used. The software *R* is used to call the Stan program via the *RStan* package [39] and analyse the MCMC output. Each simulation uses 4 MCMC chains, each with 2500 iterations and a burn–in of 1000 iterations. MCMC convergence for each parameter is checked visually using the trace plot and numerically by maintaining a 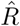 statistic of at most 1.1 [40, 41]. The 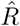 statistic is the ratio of between–chain variance to within–chain variance. All post warm–up samples are used to compute posterior distributions. To investigate differences between primary and secondary groups in the estimated values of the patient–specific parameter, *V*_0_, the joint posterior distributions across all patients within a group are compared.

In order to assess how well the two model variants capture patient and group–level variations in virus dynamics, we use posterior predictive plots and the widely applicable information criterion (WAIC) measure [42]. The WAIC measure is commonly applied for estimating expected predictive accuracy from a fitted Bayesian model using the log–likelihood evaluated at the posterior simulations of the parameter values, and is asymptotically equivalent to leave–one–out cross validation [43]. Following the description in [43], we compute the WAIC measure using the log pointwise predictive density (*lpd*) and the estimated effective number of parameters (*P*_*eff*_) as:

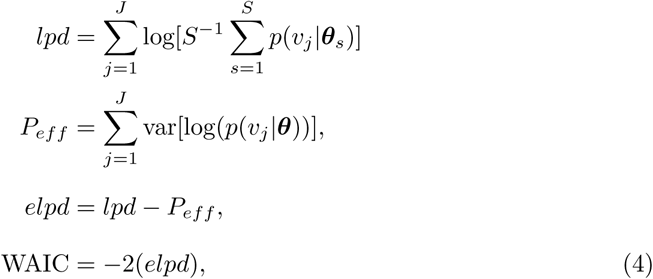

where *J* is the number of patients, *S* is the number of posterior samples, ***θ*** are the parameters, and *v* is the data [43]. The term (*lpd* − *P*_*eff*_) is the expected log pointwise predictive density (*elpd*). We estimate *P*_*eff*_ using the posterior variance of the log predictive density for each patient data, *v*_*j*_. That is, *V*_*s*_^*S*^ = log(*p*(*y*_*j*_|***θ***_*s*_)), where *V*_*s*_^*S*^ stands for the sample variance, which is given as 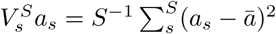. Finally, we sum over all patients to estimate the effective number of parameters. To assess the uncertainty of the WAIC estimate of prediction error, we compute its standard error using the individual components of *elpd* as:

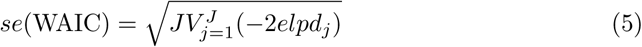

To compare the two fitted models, we estimate the difference in their predictive accuracy by the difference in *elpd*. The standard error of this difference is computed as:

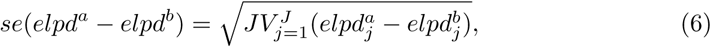

where *elpd*^*a*^ and *elpd*^*b*^ are the *elpd* measures of model *a* and model *b*, respectively.

We choose the WAIC measure over the deviance information criterion (DIC) because the DIC has the limitation of relying on a point estimate rather than the full posterior distribution. An alternative approach to compare the two models is to use the Bayes factor (BF), which is the ratio of the marginal likelihoods of each model. However, in general, the marginal likelihoods and BFs can be computationally difficult to estimate accurately and may be sensitive to prior specifications, so we avoid computing the BF. For a detailed mathematical description of DIC and BF, we direct the interested reader to [40].

### Effect of DENV–TIPs therapy

We use our ODE model (1) to assess the impact of treating DENV1 infected patients with DENV–TIPs. Previous statistical analysis of the data which we use in this study showed that increases in a patient’s plasma viremia level as one of the factor associated with higher probability of DENV transmission from humans to mosquitoes, and that the likelihood of successful transmission coincides with the period at which patients show clinical symptoms, with few transmissions occuring after the day of defervescence [15]. Thus, our goal here is to determine if administering excess DENV–TIPs to patients can reduce peak viremia and/or the duration of infection, and if so, the smallest DENV–TIPs dose required for an effect.

The effect of DENV–TIPs therapy is examined using two simulation studies. The first study concerns administering different doses of DENV–TIPs and varying the effectiveness parameter, *σ*, to an untreated DENV1 infected patient on fever day −4 which is assumed to be the time at which disease symptoms start to appear. The aim of this experiment is to determine if treatment depends on the effectiveness of DENV–TIPs as well as the concentration of DENV–TIPs dose used. The posterior medians for each estimated parameter from model *a* are used to simulate the DENV1 profile of an untreated patient, We expect that this profile represents the dynamics of a *typical* DENV1 infected patient in the data. We discuss the reasons for using parameter estimates from model *a* in the *Results* section. Values used for *σ* are 0.5, 0.6, 0.7, 0.8, 0.9 and 1.0. These values for *σ* indicate that the effectiveness of DENV–TIPs in blocking DENV from coinfected cells are 50%, 60%, 70%, 80%, 90% and 100%, respectively. For each chosen value of *σ*, the patient is treated with a DENV–TIPs dose starting from 100 copies/ml, increased by 2 fold on log10 scale until 10^12^ copies/ml.

In the second study, the patient is treated at two time points before DENV1 peaks (fever days −8 and −6) and two time points after the peak (fever days −2 and 0), testing with the same *σ* values as used in the first simulation study. The intuition behind treatment before DENV1 peak is that patients usually seek healthcare after the onset of symptoms, at which time DENV1 level is around the peak, and whether treatment may be targeted to those who do not show symptoms but have the possibility of being infected. This type of treatment may be applied to people in a close knit community such as the army or households, thereby treats all individuals in the community once the disease appears in any individual. The reason behind intervention after the peak is to investigate how effective DENV–TIPs therapy is after the onset of symptoms.

## Results

Fig 2 depicts the sensitivity of the DENV1 output from the ODE (1) to the assigned parameters (*A, ω, η, σ*). The first subplot in Fig 2 shows a baseline DENV1 trajectory (black curve in subplot (a)) which is simulated from the ODE along with measured DENV1 primary and secondary profiles (coloured lines). We consider the black curve in this subplot as a baseline, as it captures a general trend in the observed DENV1 dynamics, although there is considerable variability in the observed dynamics. To compute the baseline curve, we first used the values given in Table 1 for the assigned parameters, and the values for estimated parameters as: *β* = 3.83 × 10^−11^ (or *β* = 5 × 10^−11^), *κ* = 3.5, *δ* = 3.3, *γ* = 2, *µ* = 1 × 10^−4^ and *V*_0_ = 1. The values for *β, κ, δ* and *V*_0_ are taken from the fit of the virus neutralisation model of [31] to their DENV1 data. The value for *µ* is taken from the within–host dengue model studied in [35]. We envisaged that the death rate of coinfected cells, *γ*, would be less than the death rate of cells infected with only DENV1, *δ*, and so assigned for *γ* a value less than *δ*. With these values, however, the computed DENV1 trajectory was substantially below the observed DENV1 profiles (not shown). Thus, we fixed values for parameters *A, σ* and *µ* as discussed above and experimented with different combination of values for the remaining parameters to obtain the baseline trajectory. The baseline trajectory (the black curve in subplots (b)–(e)) is then compared with the simulated DENV1 trajectories, after varying each assigned parameters one at a time, while keeping the remaining parameters at their baseline values. In all simulations, the initial condition used was (*S*(0), *I*_*V*_ (0), *I*_*T*_ (0), *I*_*B*_(0), *V* (0), *T* (0)) = (10^8^, 0, 0, 0, 57.122, 0), and an incubation period of 6 days was assumed.

**Fig 2.**
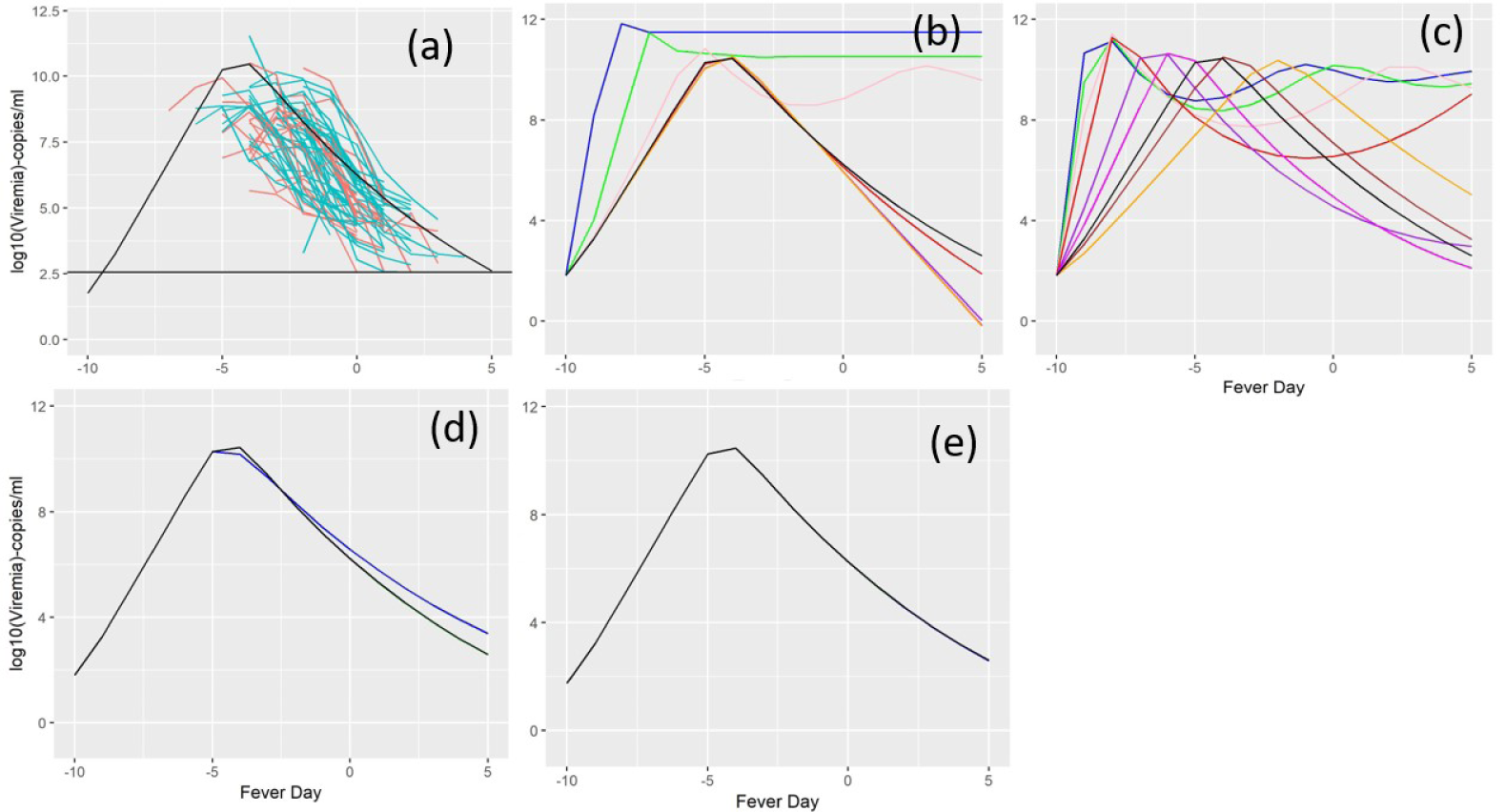
Sensitivity analysis of fixed parameters. Plot (a) shows a *baseline* DENV1 profile (black curve) computed from the ODE, with observed DENV1 primary (red) and secondary (blue) measurements. Parameter values used to compute the baseline trajectory are: *A* = 1.4 × 10^6^, *σ* = 0.5, *µ* = 0.0001, *δ* = 5.93, *γ* = 5.08, *κ* = 3.71, *η* = 42760, *ω* = 11333, *β* = 6.66 × 10^−11^. Horizontal line in plot (a) corresponds to the LOD for DENV1 data. Plots (b)–(e) show DENV1 dynamics from ODE by varying parameters *A, ω, η* and *σ*, respectively, one at a time while fixing others at their baseline value (the black curve in each plot). Values of parameter *A* used in plot (b) are: blue (10^9^), green (10^8^), pink (10^7^), black (1.4 × 10^6^), red (10^6^), purple (10^5^), magenta (10^4^), brown (10^3^), orange (100). Values of parameter *ω* used in plot (c) are: blue (10^5^), green (8 × 10^4^), pink (6 × 10^4^), red (4 × 10^4^), purple (2 × 10^4^), black (11333), magenta (1.5 × 10^4^), brown (1 × 10^4^), orange (800). Values of parameter *η* used in plot (d) are: blue (10^6^), green (1 × 10^5^), pink (8 × 10^4^), red (5 × 10^4^), purple (10^4^), black (42760), magenta (8 × 10^3^), brown (800). Values of parameter *σ* used in plot (e) are: blue (1), green (0.8), pink (0.6),red (0.4), purple (0.2), black (0.5), magenta(0).

Based on the the sensitivity plot of parameter *A* (Fig 2, subplot (b)), it appears that DENV1 output is relatively insensitive to variability in parameter *A* (susceptible cell production rate) between 100 and 1.4 × 10^6^ (baseline value), but the peak starts to occur earlier, with higher virus level at the peak for values above 1.4 × 10^6^. Moreover, as *A* is increased, the DENV1 level stabilises at a constant value after the peak. Similarly, the DENV1 peak occurs earlier for higher values of *ω* (DENV production rate), with trajectories showing waves after the peak for values above 2 × 10^4^ (Fig 2, subplot (c)). The trajectory corresponding to *ω* = 1 × 10^4^ (brown) is similar to the baseline curve (black) except for a slight shift towards the right, justifying our choice of fixing *ω* at 1 × 10^4^. The DENV1 dynamic is relatively insensitive to the values of *η* (DENV–TIPs production rate from coinfected cells) (Fig 2, subplot (d)) and *σ* (effectiveness of DENV–TIPs) (Fig 2, subplot (d)), supporting our option not to estimate these parameters.

Table 2 details the two Bayesian model variants considered, specifies which parameters were fitted as common, group– and patient– specific, and the estimated WAIC measures along with their standard errors, which were computed using Eq (4) and Eq (5), respectively. The 95% credible intervals of predicted DENV1 dynamics from model *a* are shown for five primary and five secondary patients in Fig 3 and the fits corresponding to the same patients from model *b* are given in Fig 4. Computed 95% credible intervals of DENV–TIPs and susceptible cell dynamics are also shown for each patient. The plots in Figs 3–4 does not confirm any visible distinction between the two model fits, indicating that both models can recover the observed virus profiles equally well. However, comparison of the predictive performances of the two models reveals an estimated difference in *elpd* of 3.9 (with a standard error of 1.1) in favour of model *a*, though the WAIC measures are very similar (Table 2). Hence, we present posterior predictions from model *a* fit for all patients in S1 Fig–S2 Fig. These plots suggest that the ODE (1) can recreate 49 out of 67 virus profiles reasonably well. Estimated parameter values for model *a* are shown in Table 3.

**Table 3.** Parameter estimates from model *a* fit. Parameters *δ, γ, κ, β, µ* are fitted common to all patients, and are summarised using posterior median and 95% credible interval in the curved parentheses. Initial viral load (*V*_0_) is patient–specific and is summarised by taking the median of each posterior and the median of these medians is given with IQR in the squared parentheses.

| Parameter | DENV1 Estimates |
| --- | --- |
| $\delta$ | 8.844 (6.181, 9.951) |
| $\gamma$ | 6.585 (2.970, 9.827) |
| $\kappa$ | 8.855 (6.169, 9.951) |
| $\beta \times 10^{-10}$ | 1.283 (0.9470, 1.506) |
| $\mu \times 10^{-4}$ | 5.130 (0.7348, 9.746) |
| $\sigma_e$ | 2.681 (2.443, 2.949) |
| $V_0$ | 55.29 [26.25, 94.86] |

**Fig 3.**
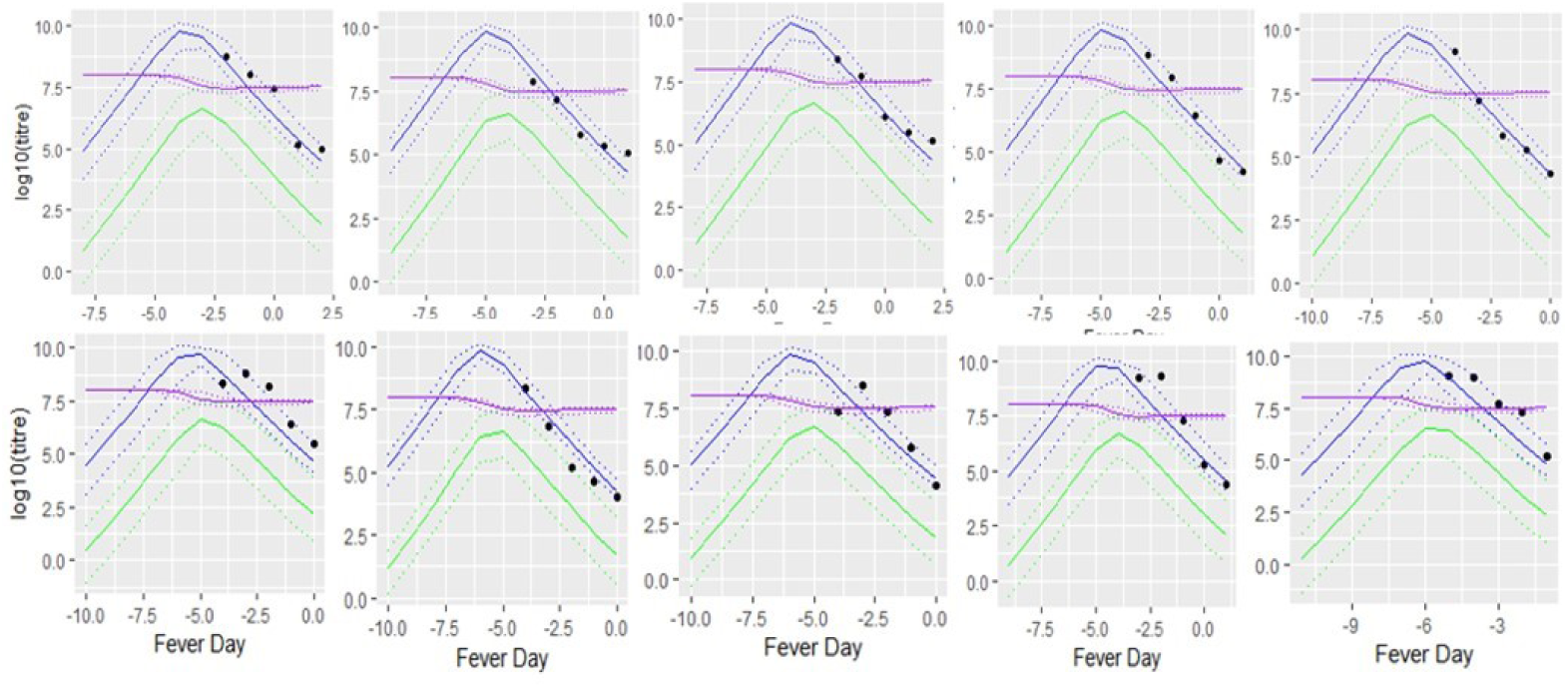
Fit of model *a* to five secondary (1st row) and five primary (2nd row) DENV1 profiles. Blue, purple and green curves represent the 95% posterior credible interval (dashed curves) for DENV1, susceptible cells and DENV–TIPs trajectories, respectively, along with the medians (solid curve), computed from all posterior samples. Measured DENV1 data are shown as black filled circles.

**Fig 4.**
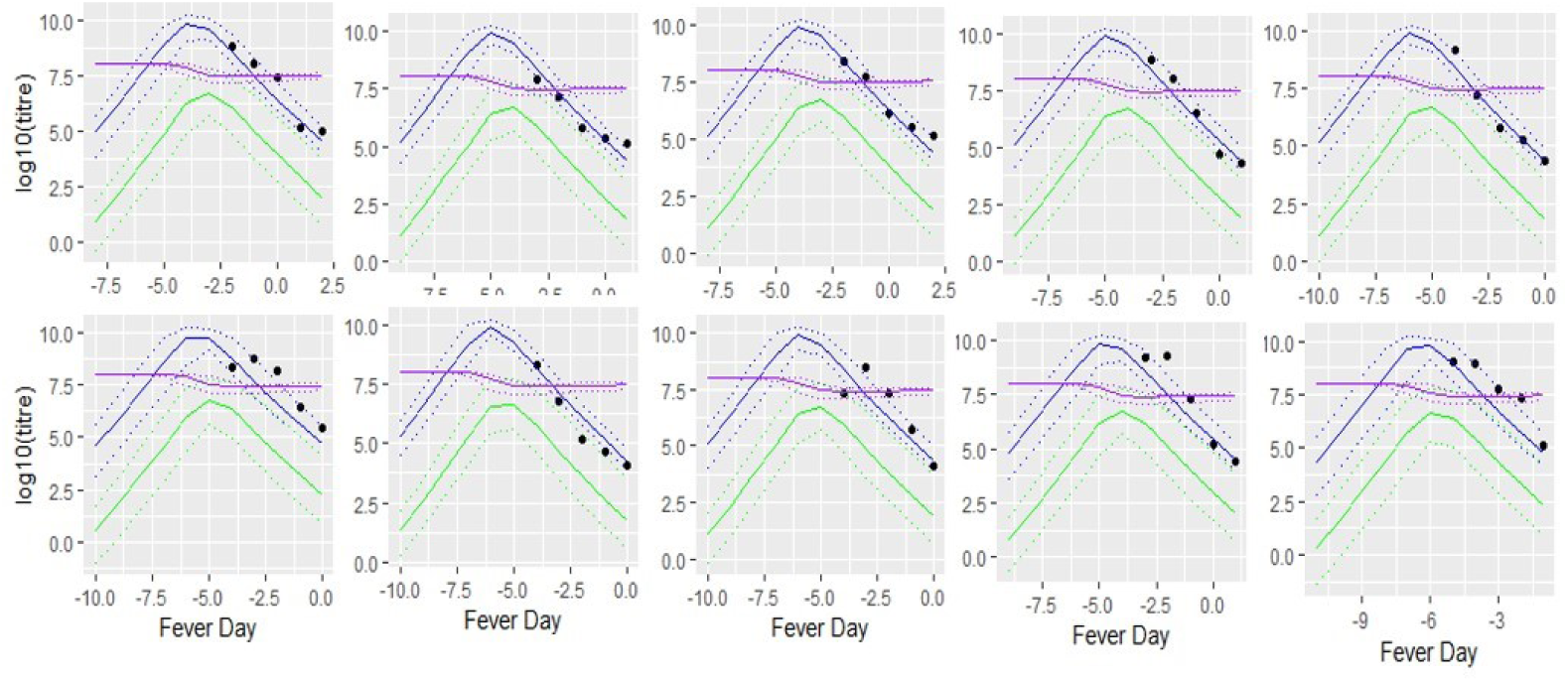
Fit of model *b* to DENV1 profiles corresponding to the same patients in Fig 3. Line types, colours and the symbol have the same meaning as given in Fig 3.

To assess potential differences between primary and secondary infections, we compare parameter estimates for these two patient groups from model *b* (Table 4), which allowed parameters *δ, γ, κ, β, µ* to vary between the two groups. These estimates indicate that there may be some chance of the two types of infection coinciding due to a large overlap in the 95% credible intervals across the group–specific parameters, although the median estimates for *δ, γ, κ* are slightly larger for the secondary infection than the primary infection. This finding agrees with the conclusion derived previously based on the *elpd* measure, that the predictive performance of model *a* is favoured over that of model *b*. On the other hand, this finding appears to contradict the theory of ADE that DENV replication (*β*) and clearance (*κ*) rates are expected to be significantly higher for secondary infection than primary infection.

**Table 4.**
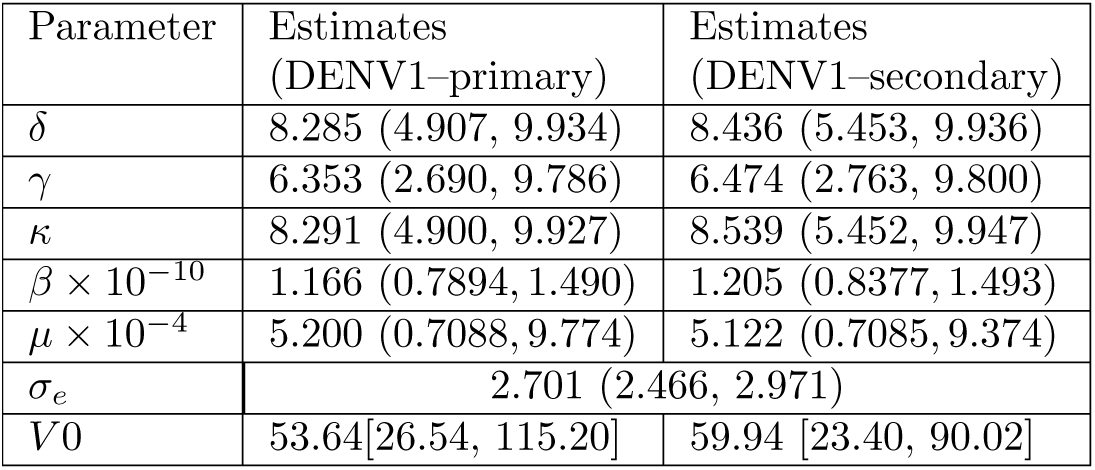
Parameter estimates from model *b* fit. Parameters *δ, γ, κ, β, µ* are fitted as group–specific, *V*_0_ is fitted as patient–specific. Parentheses have same meaning as given in Table 3. Measurement error, *σ*_*e*_ is fitted common to all patients.

We examined the impact of DENV–TIPs therapy by implementing two simulation studies. In both studies, we administered DENV–TIPs doses with varying concentration and effectiveness (modeled by parameter *σ*), to an untreated DENV1 infected patient. Our first simulation study concerns treatment of the patient on fever day −4, which is assumed to be the day of symptoms onset as it falls around a time point of DENV1 peaks (Fig 5). Our second simulation study concerns treatment of the patient on two selected days prior to symptoms onset (fever days −8 and −6, Fig 6 and Fig S3 Fig, respectively) and two selected days post symptoms onset (fever days −2 and 0, Fig 7 and Fig S4 Fig, respectively).

**Fig 5.**
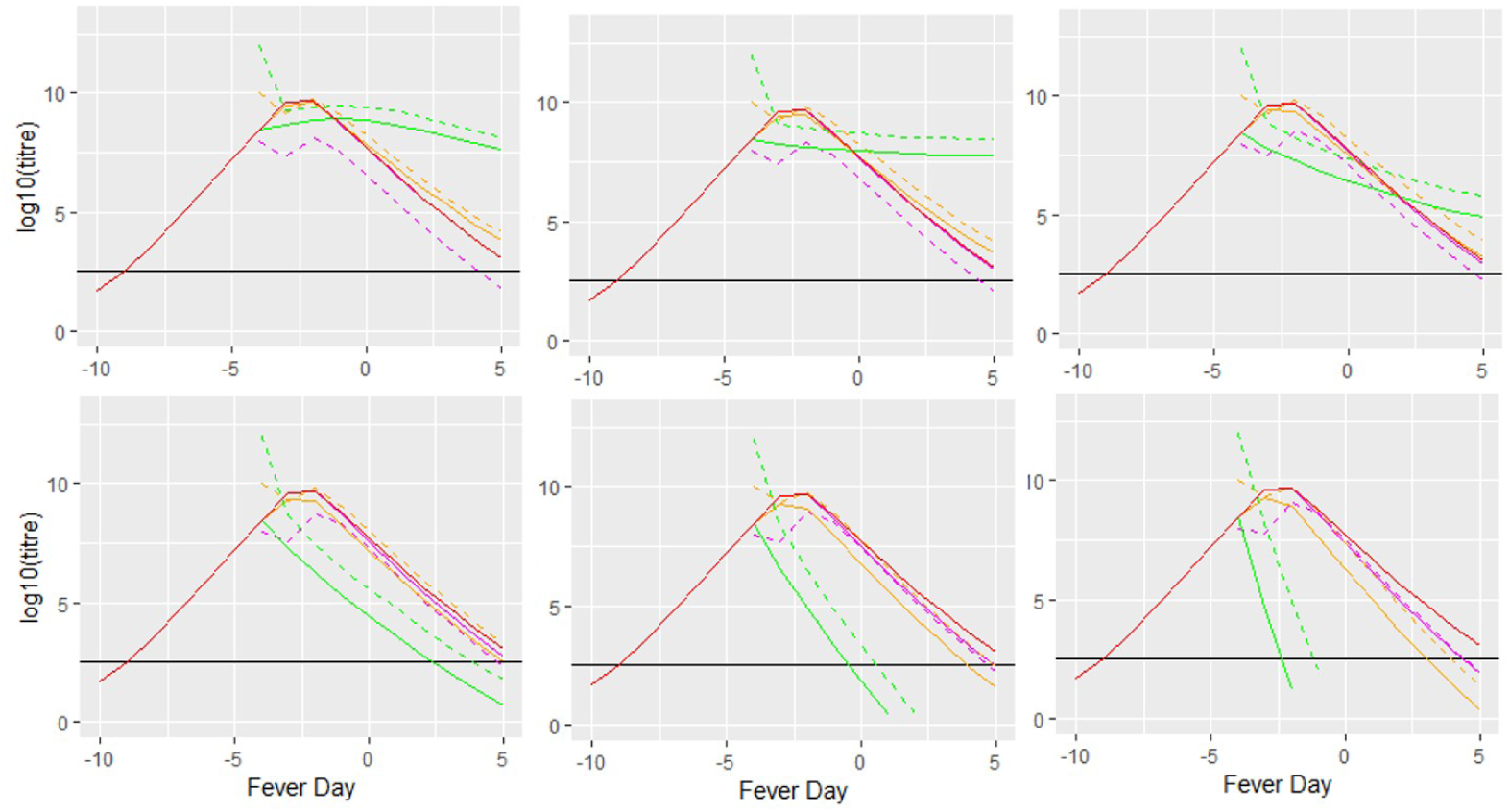
First simulation study: effect of TIPs therapy on fever day −4. Effect of treating a *typical* DENV1 patient with various concentration of DENV–TIPs doses along with varying the effectiveness parameter, *σ* on fever day −4. The solid red curve in each plot corresponds to the DENV1 profile of an untreated patient. Values of *σ* used are *σ* = 0.5 (first row, first column), *σ* = 0.6 (first row, second column), *σ* = 0.7 (first row, third column), *σ* = 0.8 (second row, first column), *σ* = 0.9 (second row, second column) and *σ* = 1.0 (second row, third column). Each coloured curve (other than the red curve) shows the profile of DENV–TIPs (dashed) and DENV1 (solid) after treatment. Concentration of DENV–TIPs (copies/ml) used are: 10^8^ (magenta), 10^10^ (orange) and 10^12^ (green). The horizontal black line corresponds to the LOD.

**Fig 6.**
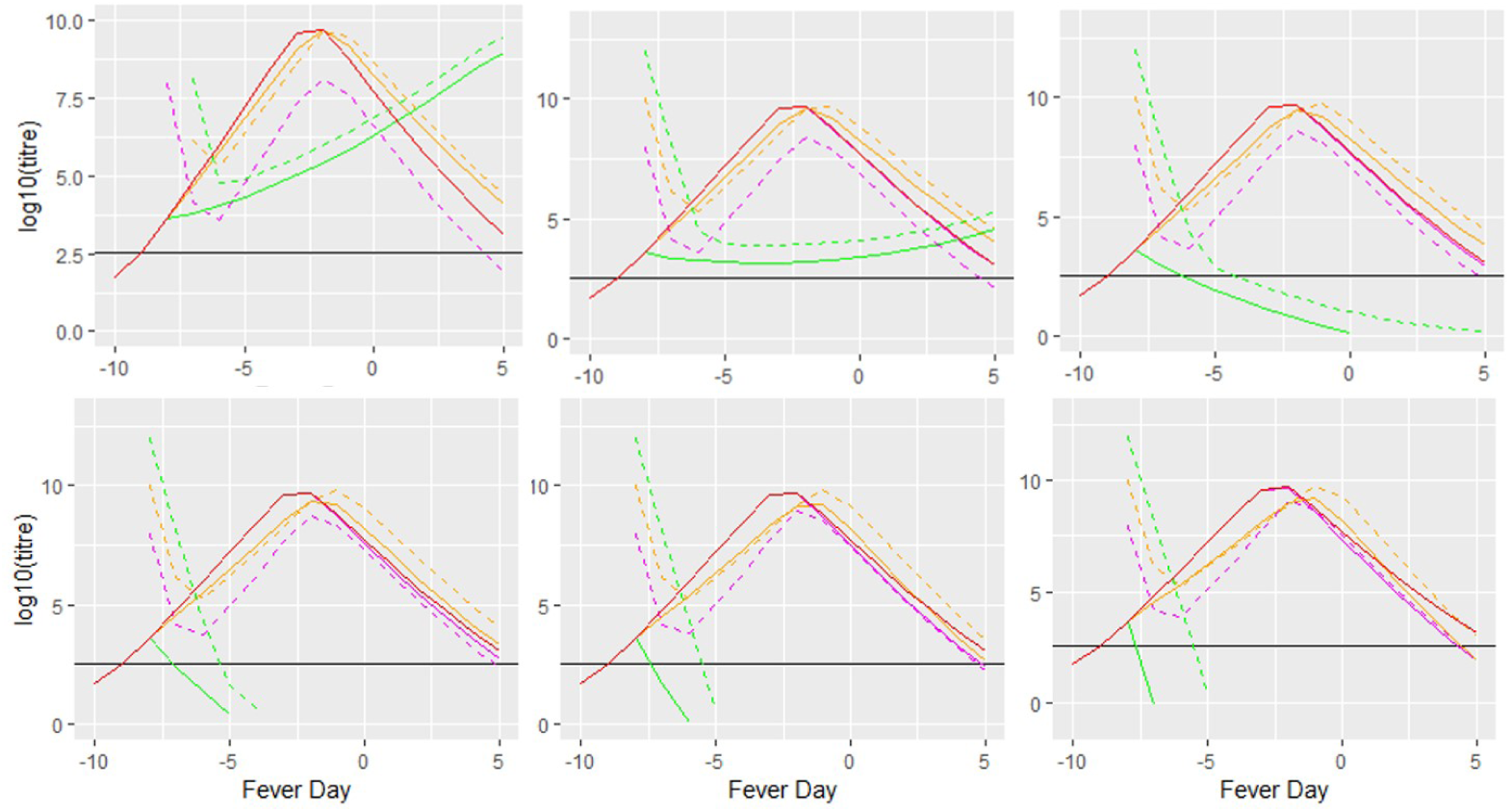
Second simulation study: effect of TIPs therapy on fever day −8. Effect of treating a *typical* DENV1 patient with various concentration of DENV–TIPs doses along with varying the effectiveness parameter, *σ* on fever day −8. The row and column position of each subplot corresponds to the *σ* value as given in Fig 5. The colours and line types have the same meanings as in Fig 5.

**Fig 7.**
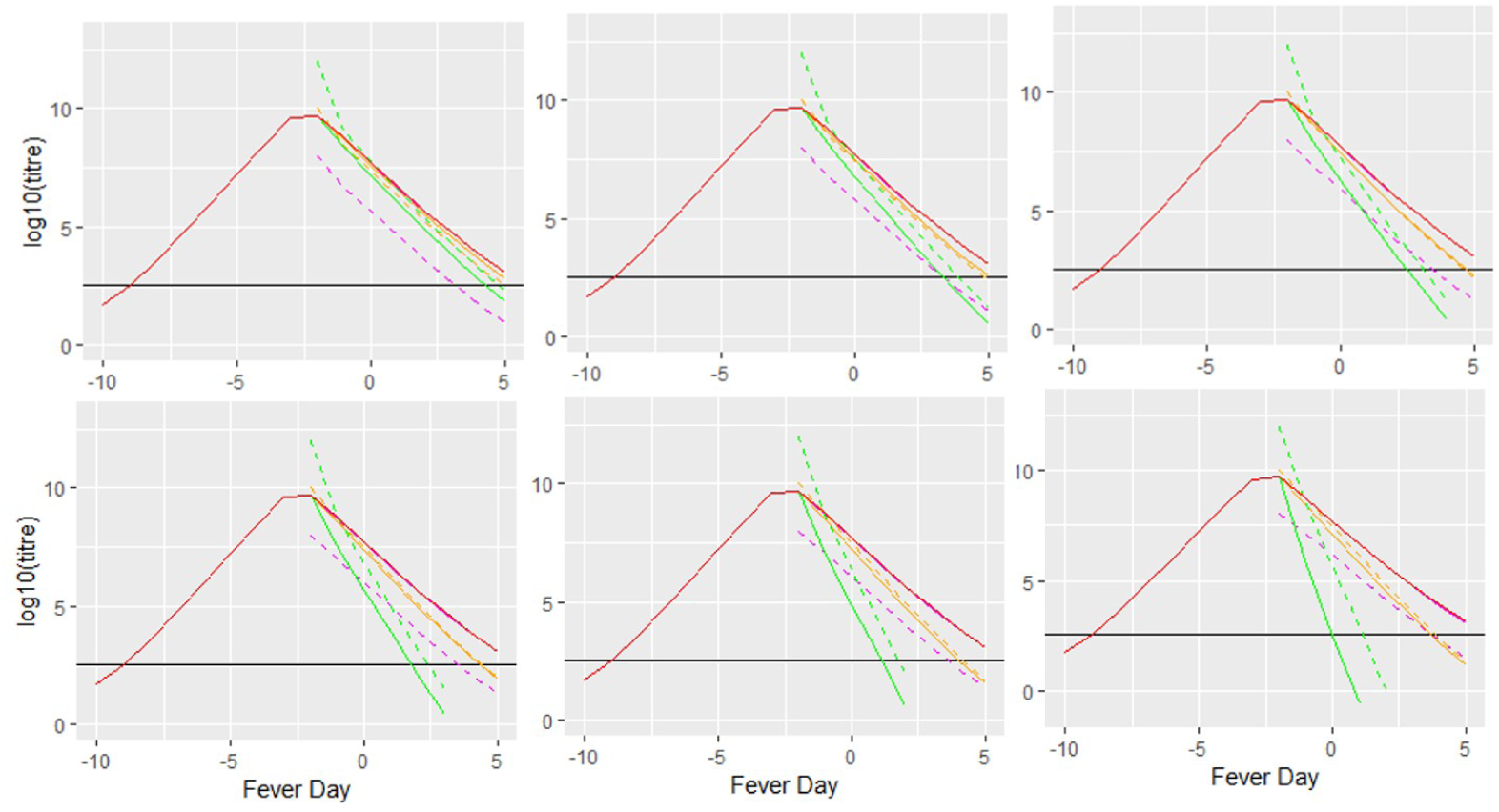
Second simulation study: effect of TIPs therapy on fever day −2. Effect of treating a *typical* DENV1 patient with various concentration of DENV–TIPs doses along with varying the effectiveness parameter, *σ* on fever day −2. The row and column position of each subplot corresponds to the *σ* value as given in Fig 5. The colours and line types have the same meanings as in Fig 5.

In both studies, we first simulated the profile of an untreated DENV1 patient (Figs 5–7 and S3 Fig–S4 Fig, the solid red trajectory in each subplot). We consider this profile to represent a general trend of an untreated DENV1 patient as it is simulated using the median summary of each estimated parameter from model *a* (Table 3), which is favoured over model *b*, based on the *elpd* measures. We then administered DENV–TIPs doses to this patient, starting from 100 copies/ml, increasing by 2 fold on log10 scale, until 10^12^ copies/ml, by varying effectiveness parameter, *σ*.

Our first simulation study shows that the impact of DENV–TIPs treatment, at the time of symptoms onset, on DENV1 dynamics depends on both the effectiveness parameter, *σ*, and the concentration of DENV–TIPs dose administered (Fig 5). In general, a dose lower than 10^8^ copies/ml of DENV–TIPs (not shown) does not have any discernible affect on the DENV1 dynamics for any chosen values of *σ*. When *σ* is at most 0.7 and the dose administered is above 10^8^ copies/ml, the DENV1 peak starts to decrease, but the duration of infection also increases (Fig 5, the three subplots in the first row, orange and green solid curves). On the other hand, when *σ* is at least 0.8, both duration and peak start to decrease for doses at and above 10^8^ (Fig 5, the three subplots in the second row, magenta, orange and green solid curves), with a sharp reduction in duration for the dose with 10^12^ copies/ml (Fig 5, second row, green solid curve), and reaching the LOD within about 4 and 2 days when *σ* is 0.9 and 1.0, respectively.

The second simulation study shows that, when treatment is target prior to symptom onset, a sharp reduction in duration of infection occurs for the dose with 10^12^ copies/ml, if *σ* is at least 0.7 (Fig 6 and Fig S3 Fig, subplots starting from first row third column to the last subplot, the green solid curve), If *σ* is less than 0.7 and the dose administered is 10^12^ copies/ml, DENV1 level at peak is reduced, but duration of infection increases (Fig 6 and Fig S3 Fig, the first and second subplots in the first row, the green solid curve). Similarly, If *σ* is at most 0.7 and the dose administered is 10^10^ copies/ml, duration of infection increases slightly (Fig 6 and Fig S3 Fig, the three subplots in the first row, the orange solid curve). A dose of 10^8^ copies/ml has negligible impact on DENV1 profile (Fig 6 and Fig S3 Fig, the magenta solid curve in each subplot).

Similar to the results obtained for treatment prior to symptoms onset, the most effective dose for treatment after symptoms onset is 10^12^ copies/ml, but with *σ* being at least 0.6 (Fig 7 and Fig S4 Fig, subplots starting from first row second column to the last subplot, the green solid curve). A dose of 10^10^ copies/ml given after symptom onset also reduces duration of infection (Fig 7 and Fig S4 Fig, subplots starting from first row second column to the last subplot, the orange solid curve), but to a lesser extent than that for the 10^12^ copies/ml dose. A dose of 10^8^ copies/ml has no visible impact on DENV1 profile (Fig 7 and Fig S4 Fig, the magenta solid curve in each subplot).

Both simulation studies reveal that a dose of 10^8^ copies/ml has negligible impact on DENV1 profile at any of the selected time points and for any chosen *σ* value (Figs 5–7 and Figs S3 Fig–S4 Fig, the magenta solid lines in each subplot) and lower (not shown). They also show that the most effective dose of DENV–TIPs for intervention of DENV1 infection at any time point of infection is 10^12^ copies/ml. However, the effectiveness of DENV–TIPs in blocking DENV1 from coinfected cells (modeled by parameter *σ*) must be at least 0.8, 0, 7 and 0.6 for treatment targeted at the time point of symptoms onset, early and late stages of infection, respectively. Thus, choosing a value of at least 0.8 for *σ*, will satisfy the condition required for effectiveness of DENV–TIPs, in order to limit disease severity at all time points during DENV1 infection, provided that the DENV–TIPs dose administered is 10^12^ copies/ml.

## Discussion

We introduced a system of an ordinary differential equations (ODE) model for the dynamics of dengue wild–type virus (DENV) transmissible interfering dengue particles (DENV–TIPs) as they interact with human host cells, incorporating effectiveness of DENV–TIPs in blocking DENV from coinfected cells. Parameters of the ODE model were estimated by fitting it to DENV1 infected patient data. By allowing the initial concentration of DENV1 to vary we were able to recover the observed variation between patients. To the knowledge of the authours, this is the first study in which effectiveness of DENV–TIPs in limiting DENV replication is modelled and validated using patient data.

We examined the sensitivity of the assigned parameters to DENV1 output from the ODE model. We found that the DENV1 output was relatively insensitive to variability in the effectiveness parameter (*σ*) and the rate of production of DENV–TIPs from coinfected cells (*η*). For the susceptible cells production rate (*A*), values larger than 1.4 × 10^6^ produced DENV1 trajectories that did not resemble observed data, while keeping DENV1 production rate (*ω*) at the assigned value 10^4^. Similarly, unreasonable DENV1 outputs were seen for values of *ω* at and above 4 × 10^4^, while *A* was fixed at 1.4 × 10^6^.

We did not find any substantive differences between primary and secondary DENV1 infection groups. This finding does not lend support to the theory of ADE, which proposes that antibodies produced in a primary infection aid virus entrance to cells in a secondary heterologous infection as they cannot completely neutralise virus present in the latter infection, thus increasing virus infectivity rate (*β*) for secondary infection compared to primary infection. The ADE phenomenon was supported in previous within–host dengue modelling studies [29–32]. Similarly, studies [29, 32] established that the virus clearance rate, *κ*, was significantly higher for secondary infection than that of primary infection, due to an increased activation of the immune system that occurs in secondary infection.

We note that our modelling approach is rather different from the approaches used in [29–32]. These studies aimed to assess the roles of either adaptive immunity via antibodies [31, 32] or both innate and adaptive immunity via Type 1 interferon and T–cells [29, 30], in shaping DENV dynamics, without considering a role of DENV–TIPs in this process. On the other hand, we focused on investigating the role that DENV–TIPs play in shaping DENV profiles, neglecting the role which immunity may play in the process. There are also some differences between the data analysed in our study and those used in these previous studies. Patients in their study were recruited within 48 hours of fever onset, whilst patients in our study we used were recruited within 72 hours of fever onset. Thus, DENV measurements around the peak were available for many patients in their data while this information is not available for many patients in the data we used. For example, in the [31] study, 38% of patients have DENV1 measurements available around peak (S5 Fig, filled circles), but in our data only 16% of patients have DENV1 measurements around the peak. Furthermore, compared to only four to five DENV1 measurements per patient available in our data, each patient in the [31] study has about 13 DENV1 measurements. Thus, with more data points per patient and measurements about peak viremia available for many patients, their model fits may have may have been able to more confidently evaluate the above mentioned ADE phenomenon between primary and secondary infections.

In order to examine if the difference in results between our study and that of [31] may be due to the differences in the two data sets, our ODE model was fitted separately to the DENV1 primary and secondary data in [31] (S5 Fig). However, the results showed that there is a chance that the two types of infections coincide, due to the overlap in the credible intervals for parameters *β, δ, γ, κ* and *µ*, across the two groups (S1 Table). Thus, the differences in results concerning the primary and secondary groups in [31] and our analysis may not be due to the differences seen in the two data sets, but may be due to the two mathematical modelling approaches used in the two analyses. Hence, results obtained in [29–32] and ours must be interpreted with regard to the context of application.

Modelling the dynamics of both DENV–TIPs and immunity may reveal any biological differences between primary and secondary infections. The model developed in [44] accounts for the roles of both DENV–TIPs and antibodies, and may be used in this regard. However, unlike our model, their model does not account for effectiveness of DENV–TIPs in blocking DENV production from coinfected cells. Since antibodies usually arise late in infection, especially in primary infection, modelling the dynamics of antibodies along with DENV–TIPs may not fully highlight the role of DENV–TIPs in shaping DENV replication. Hence, we did not consider the dynamics of antibodies in our model. The study in [44] also estimated parameters of their model, but unlike our method, they implemented a Latin Hypercube Sampling method whereby parameters were uniquely sampled from a multi–dimensional grid, and the samples were used to simulate virus trajectories from their model and selected the candidate trajectories which best represented the observed data.

We assessed the impact of DENV–TIPs therapy on peak DENV1 level and the duration of infection by administering DENV–TIPs doses to an untreated DENV1 patient, with varying the concentration and effectiveness parameter (*σ*), as well as varying the time points during infection at which the treatment was given. We found that, when treatment was targeted at the time of onset of symptoms, the DENV1 level quickly reached to the limit of detection within about 2 to 4 days after the onset of symptoms when the value of *σ* was kept at least 0.8 and the dose administered was 10^12^ copies/ml. However, when the same dose was administered with values of *σ* kept below 0.8, the peak DENV1 level reduced about 2 fold on the log10 scale, but the duration of infection was increased. DENV–TIPs doses less than or equal to 10^10^ either had no effect on the DENV1 profile, or reduced the peak and duration slightly. A dose as high as 10^12^ copies/ml was also required to reduce both the duration of infection and peak DENV1 level, when treatment was targeted prior to the onset of symptoms or later stages of infection, provided that *σ* was kept at least 0.7 and 0.6, respectively. These findings reveal that choosing a value of at least 0.8 for *σ* would be sufficient in intervention of DENV1 infection at any stages of infection, provided that the DENV–TIPs dose administered is 10^12^ copies/ml.

We note that our method of assessing the impact of DENV–TIPs therapy is different from the anti–viral treatment applied in [32, 44]. In both of these studies, the treatment was specific to each patient, whilst in our approach, it was applied to a patient whose infection dynamics may represent an average patient from the population, as we used the median estimates of the posterior summary to simulate the DENV1 of this patient. In [32], similar to our method, treatment was targeted at the time of onset of symptoms, but, in general, the impact on secondary DENV1 cases was predicted to be less than on primary DENV1 cases. Unlike this result, our estimated dose was equally effective for both primary and secondary infections. In [44], various DENV–TIPs doses were administered continuously to each patient until the DENV was cleared from the body. For many DENV1 infected patients, it was required that doses as high as the DENV1 level at peak needed to be administered, for the entire duration of infection. In contrast to this finding, our result showed that a single DENV–TIPs dose of 10^12^ copies/ml, administered on any day was sufficient to reduce severity of dengue for DENV1 patients.

To compare our method of assessing DENV–TIPs therapy with that applied in [32], we investigated the impact of administering DENV–TIPs doses to ten DENV1 infected patients, by varying the effectiveness parameter, *σ*, on the reported day of symptoms (S6 Fig–S9 Fig), four days prior to the reported day of symptoms (S10 Fig–S12 Fig) and two days after the reported day of symptoms (S13 Fig–S16 Fig). For each patient, the impact of DENV–TIPs therapy on the posterior median DENV1 profile from model *a* fit was used to perform the investigation (S6 Fig–S16 Fig, the blue curve in each subplot). This investigation revealed that, similar to our conclusion from the first and second simulation studies, the most effective DENV-TIPs dose in reducing duration of infection at all chosen time points was 10^12^ copies/ml, provided that *σ* was kept at least 0.6, 0.8 and 0.7 when treatment was targeted on the day of reported symptoms onset, four days prior to the reported day of symptoms onset, two days after the reported day of symptoms onset, respectively (S6 Fig–S16 Fig, the green curve in each subplot). A dose of 10^8^ or 10^10^ does not show any visible impact on the DENV1 profile of any patient at any selected time points (S6 Fig–S16 Fig, the magenta and orange curves, respectively, in each subplot). Thus, choosing a value of at least 0.8 for *σ* would be sufficient in reducing severity of DENV1 infection at all time points, provided that the DENV–TIPs dose administered was 10^12^ copies/ml. This provides the same conclusion that we obtained from our first and second simulation studies.

A shortcoming of our analysis is that we assumed that the effectiveness of DENV–TIPs in blocking DENV from coinfected cells is the same, regardless of the amount of DENV–TIPs and DENV present in the body (that is assigning a value from 0 to 1 for parameter *σ*). As a result, DENV production rate from coinfected cells (*ω*) is scaled by a fixed amount. Ideally, DENV production rate from coinfected cells will depend on the concentration of DENV–TIPs in these cells. This amounts to assuming that if the rate of production of DENV from coinfected cells increases then the rate of production of DENV–TIPs will be reduced and vice versa. These two assumptions may be incorporated in the model by the use of monotonically decreasing and increasing hill functions [45] which depends on the concentration of DENV–TIPs in the body.

Although we did not incorporate any immune response in our ODE model, previous within–host dengue studies showed that immunity plays a significant role in shaping patients’ DENV profiles [29–32]. Thus, modelling immunity along with DENV–TIPs may reveal possible differences between primary and secondary infections, which we are not able to establish here.

Our results indicated some role for susceptible cell depletion around peak DENV level, which could be due to the fact that the model fits are calibrated to cope with the absence of data on DENV–TIPs and the four types of cells considered. Measurements of these components would help to confirm the predicted dynamics.

## Conclusion

We fitted an ODE model to DENV1 primary and secondary data, accounting for effectiveness of DENV–TIPs play in shaping patients’ DENV1 profiles, with the resulting parameter estimates confirming that variability in initial DENV1 load is sufficient to recreate patient variability in DENV1 dynamics, and not showing parameter differences between primary and secondary infections. In addition, we explored the possible impact of DENV–TIPs therapy and found that a dose as high as 10^12^ copies/ml of plasma is required to reduce the severity of DENV1 infection when the effectiveness of DENV–TIPs is at least 80%. This finding should be of use in the development of testing of DENV–TIPs for patients. Detailed data on DENV–TIPs and cell types titre would allow more comprehensive models to be developed. This work could be extended to analyse DENV data on different serotypes or on data which incorporate more patient categories such as dengue fever, dengue hemorrhagic fever and dengue shock syndrome. Other extensions of this work could be to improve the mathematical model by incorporating a component of immunity and scaling the rates of production of DENV and DENV–TIPs from coinfected cells.

## Supporting information

**S1 Fig.**
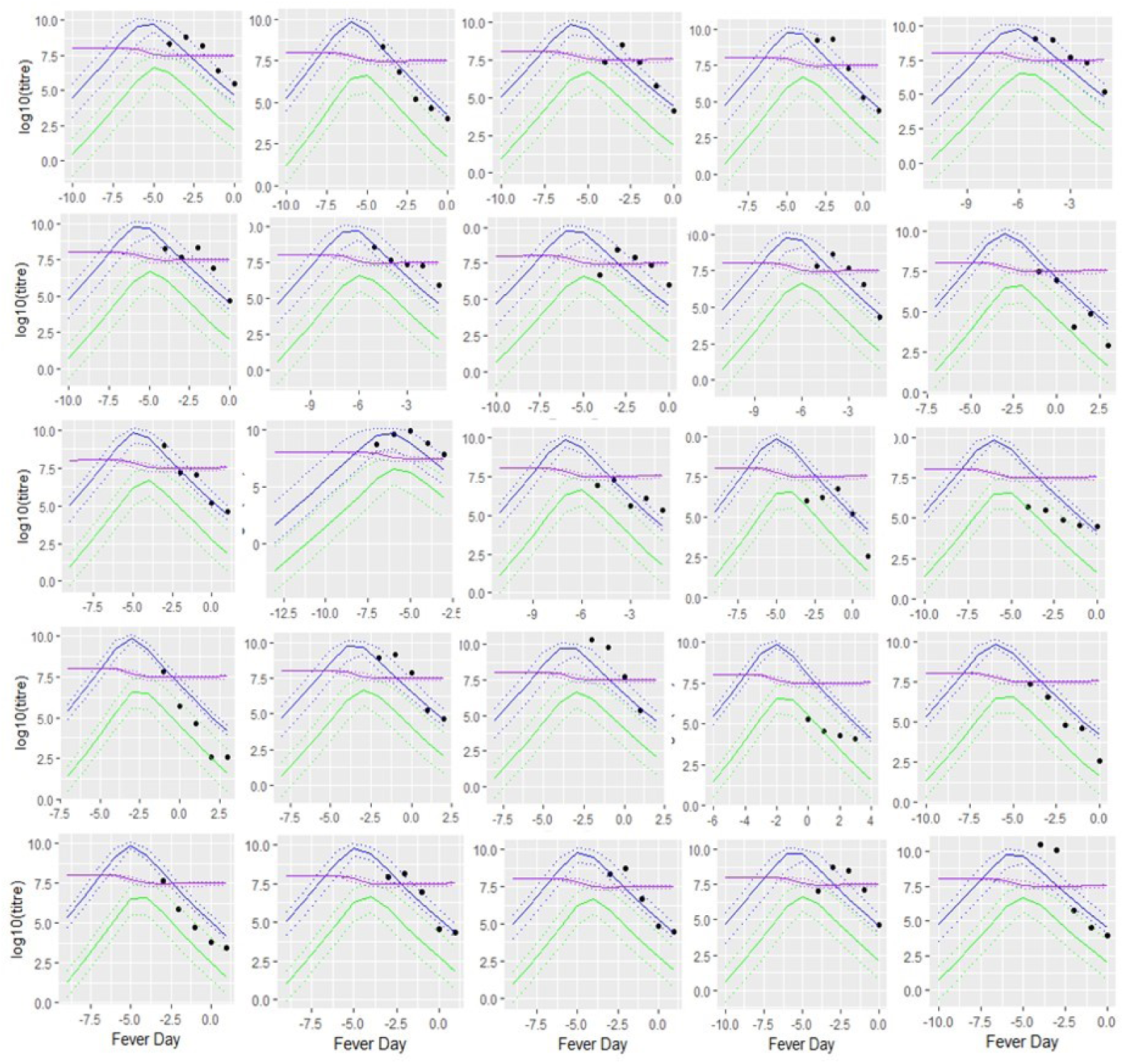
Posterior predictive plots of DENV1 primary patient data for model *a* fit. Blue, purple and green curves represent the 95% posterior credible interval (dashed) along with the median (solid) for DENV1, susceptible cells, and DENV-TIPs, respectively, computed from all posterior samples. Filled circles represent measured DENV1 data.

**S2 Fig.**
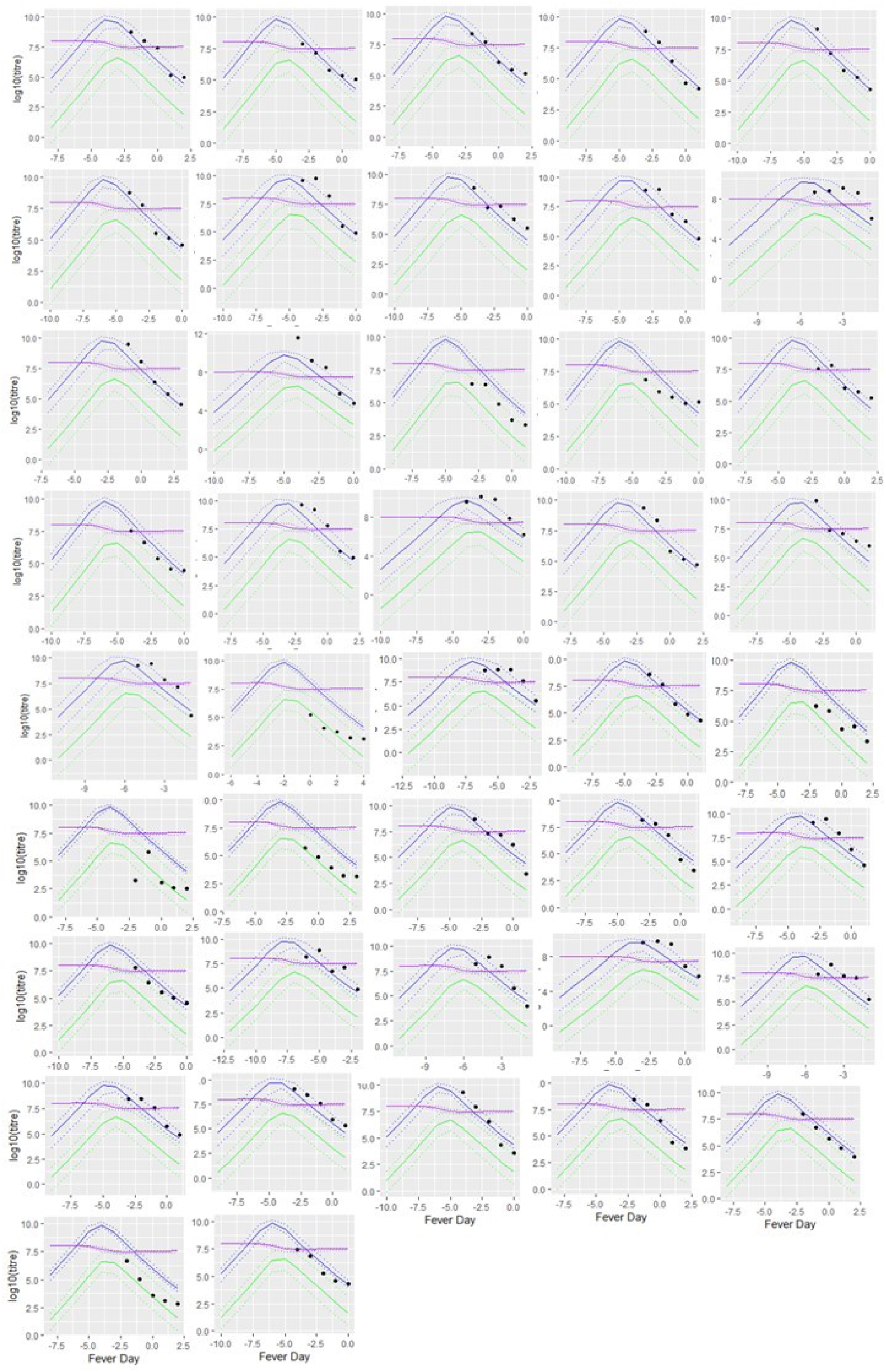
Posterior predictive plots of DENV1 secondary patient data for model *a* fit. Line types, colours and symbol have the same meaning as described in S1 Fig.

**S3 Fig.**
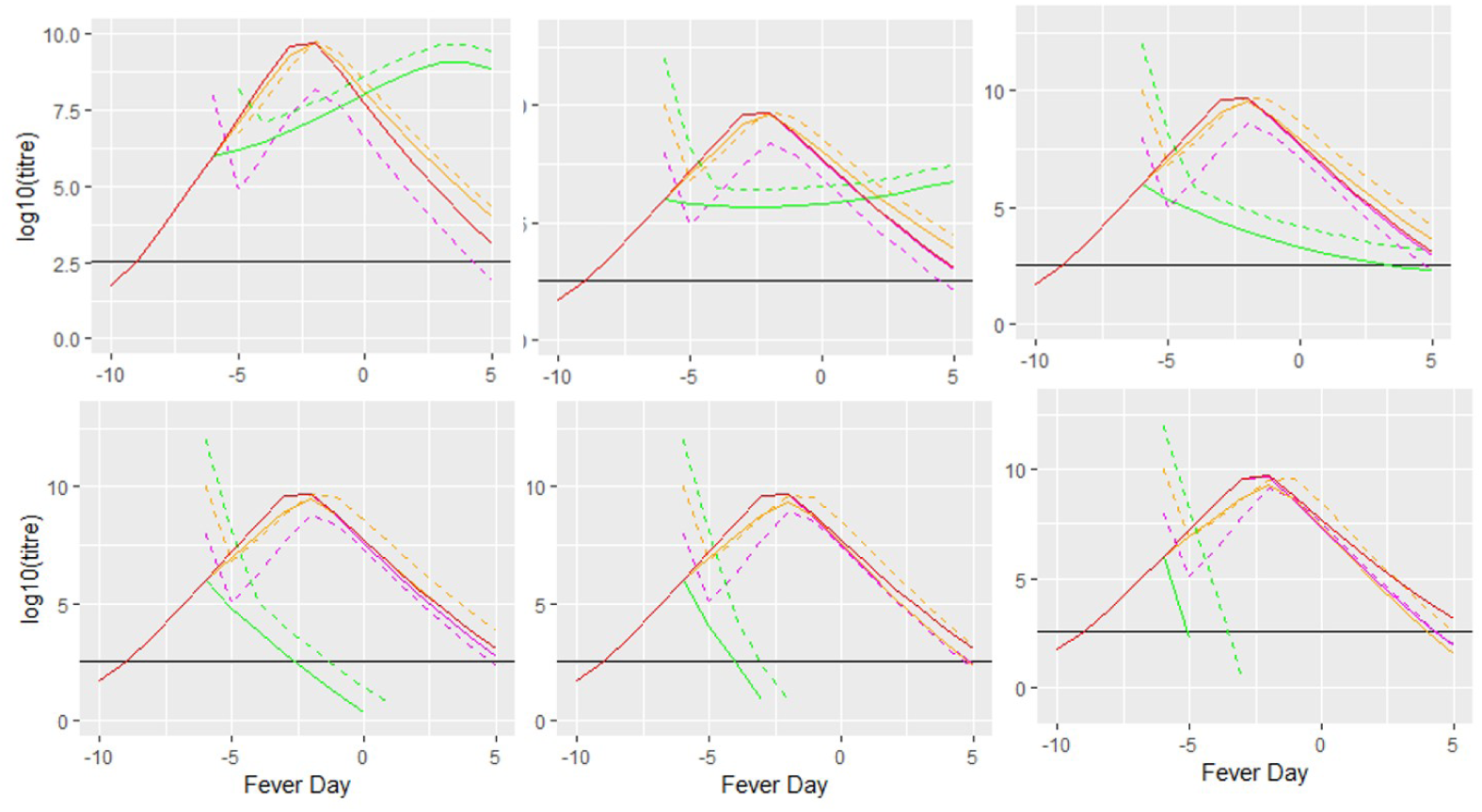
Second simulation study: effect of TIPs therapy on fever day −6. Effect of treating a *typical* DENV1 patient with various concentration of DENV–TIPs doses along with varying the effectiveness parameter, *σ* on fever day −6. The solid red curve in each plot corresponds to the DENV1 profile of an untreated patient. Values of *σ* used are *σ* = 0.5 (first row, first column), *σ* = 0.6 (first row, second column), *σ* = 0.7 (first row, third column), *σ* = 0.8 (second row, first column), *σ* = 0.9 (second row, second column) and *σ* = 1.0 (second row, third column). Each coloured curve (other than the red curve) shows the profile of DENV–TIPs (dashed) and WT–virus (solid). Concentration of DENV–TIPs (copies/ml) used are: 10^8^ (magenta), 10^10^ (orange) and 10^12^ (green). The horizontal black line corresponds to the LOD.

**S4 Fig.**
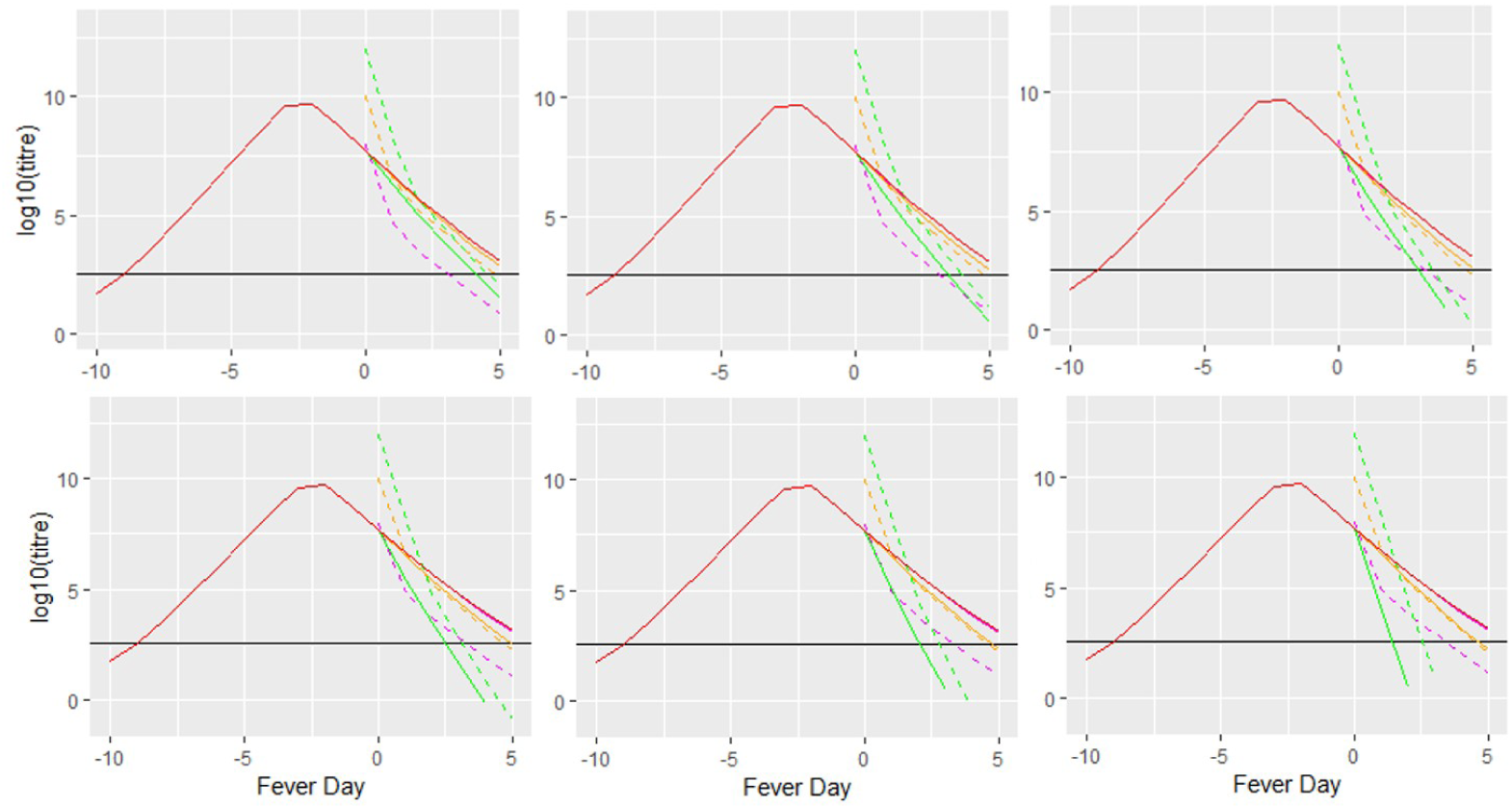
Second simulation study: effect of TIPs therapy on fever day 0. Effect of treating a *typical* DENV1 patient with various concentration of DENV–TIPs doses along with varying the effectiveness parameter, *σ* on fever day 0. The row and column position of each subplot corresponds to the *σ* value as given in S3 Fig. The colours and line types have the same meanings as in S3 Fig.

**S5 Fig.**
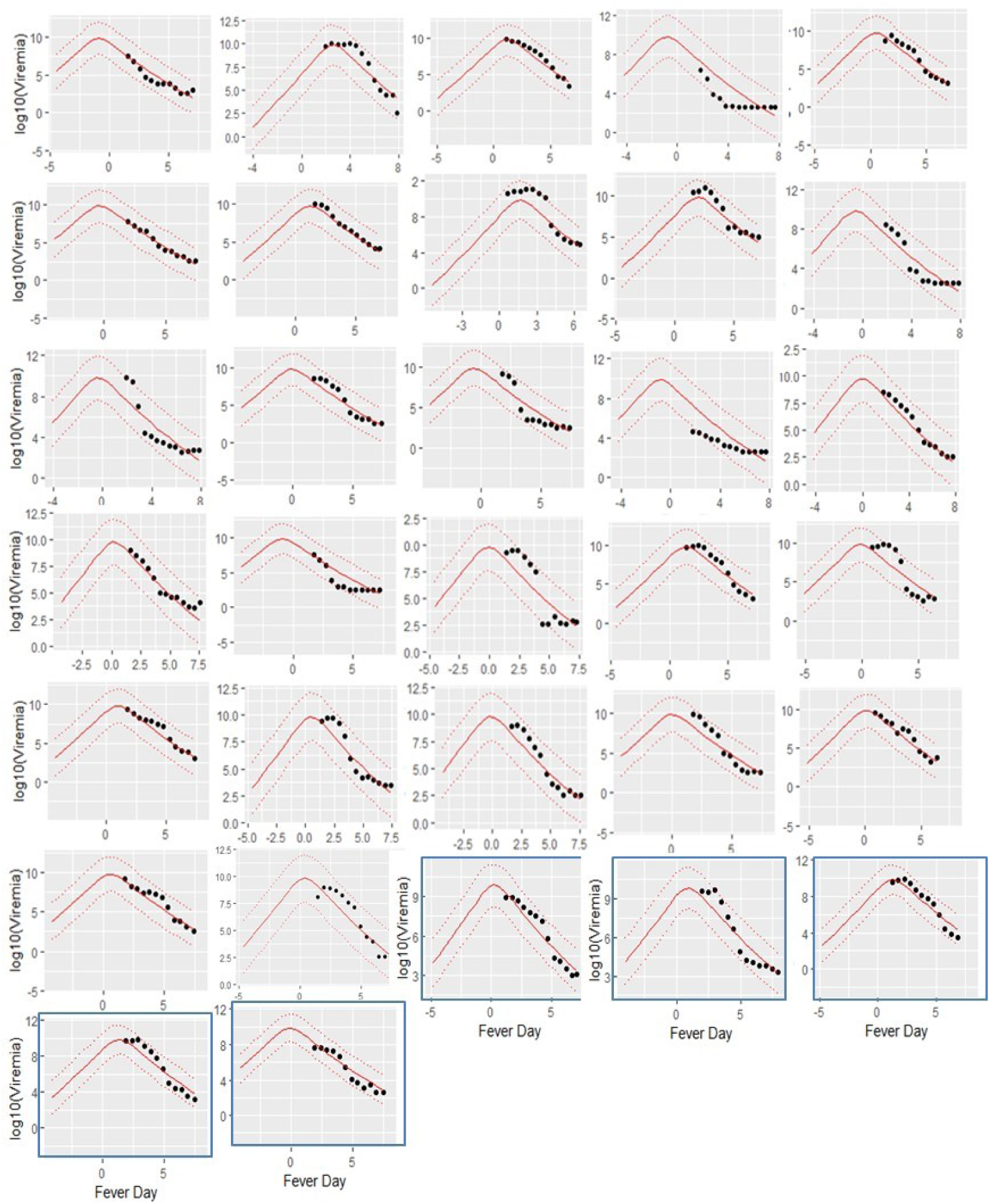
**Posterior predictive plots of DENV1 data from [31].** Model a is fitted separately to primary (n = 5, marked as blue boxes) and secondary (n = 27) DENV1 data from [31]. Dashed curves represent the 95% posterior credible interval, solid curve shows the posterior median for DENV1, computed from all posterior samples. Filled circles represent measured DENV1 data.

**S1 Table.** **Parameter estimates for the fits of model *a* separately to DENV1 primary and secondary data in [31].** Parameters *δ, γ, κ, β, µ* are summarised using posterior median and 95% credible interval in the curved parentheses. Initial viral load (*V*_0_) is patient–specific and is summarised by taking the median of each posterior and the median of these medians is given with IQR in the squared parentheses.

| Parameter | Estimates<br>(DENV1–primary, $n = 5$) | Estimates<br>(DENV1–secondary, $n=27$) |
| --- | --- | --- |
| $\delta$ | 8.473 (6.070, 9.915) | 8.972(7.441,9.935) |
| $\gamma$ | 6.787 (3.213, 9.821) | 7.122 (3.892, 9.850) |
| $\kappa$ | 8.774 (6.436, 9.949) | 9.234 (7.701, 9.965) |
| $\beta \times 10^{-10}$ | 1.247 (0.9222, 1.497) | 1.413 (1.196, 1.582) |
| $\mu \times 10^{-4}$ | 5.006 (0.6949, 9.725) | 5.003 (0.7336, 9.749) |
| $\sigma_e$ | 1.811 (1.459, 2.307) | 2.531 (2.281, 2.838) |
| $V0$ | 4.69[1.37, 6.53] | 3.44[0.3 8,42.27] |

**S6 Fig.**
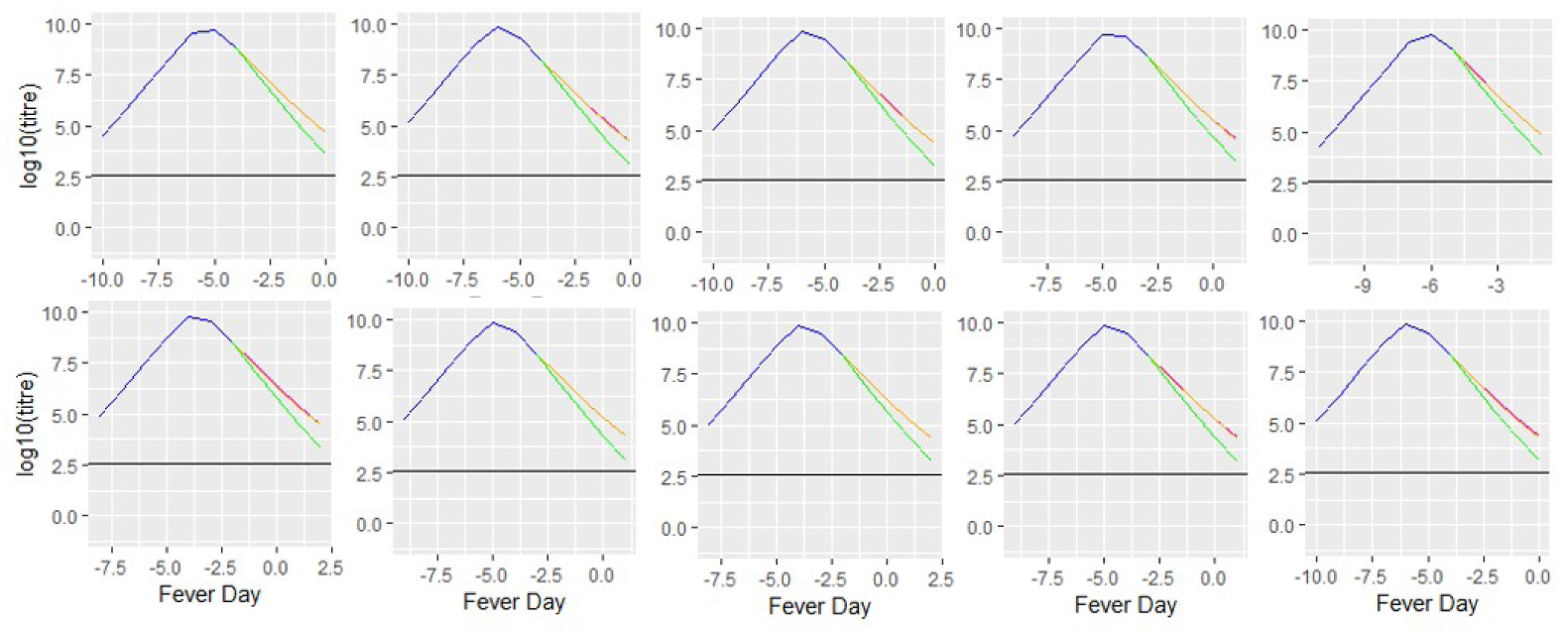
Effect of treatment of ten patients with DENV–TIPs doses on the day of reported symptoms when *σ* = 0.5. Different concentration of DENV–TIPs doses are administered to five primary (first row) and five secondary (second row) DENV1 infected patients. The blue curve in each subplot represents the posterior median for the DENV1 trajectory from model *a* fit. The magenta, orange and green curves represent DENV–TIPs doses with concentration (copies/ml) 10^8^, 10^10^ and 10^12^, respectively. Horizontal black line corresponds to the LOD.

**S7 Fig.**
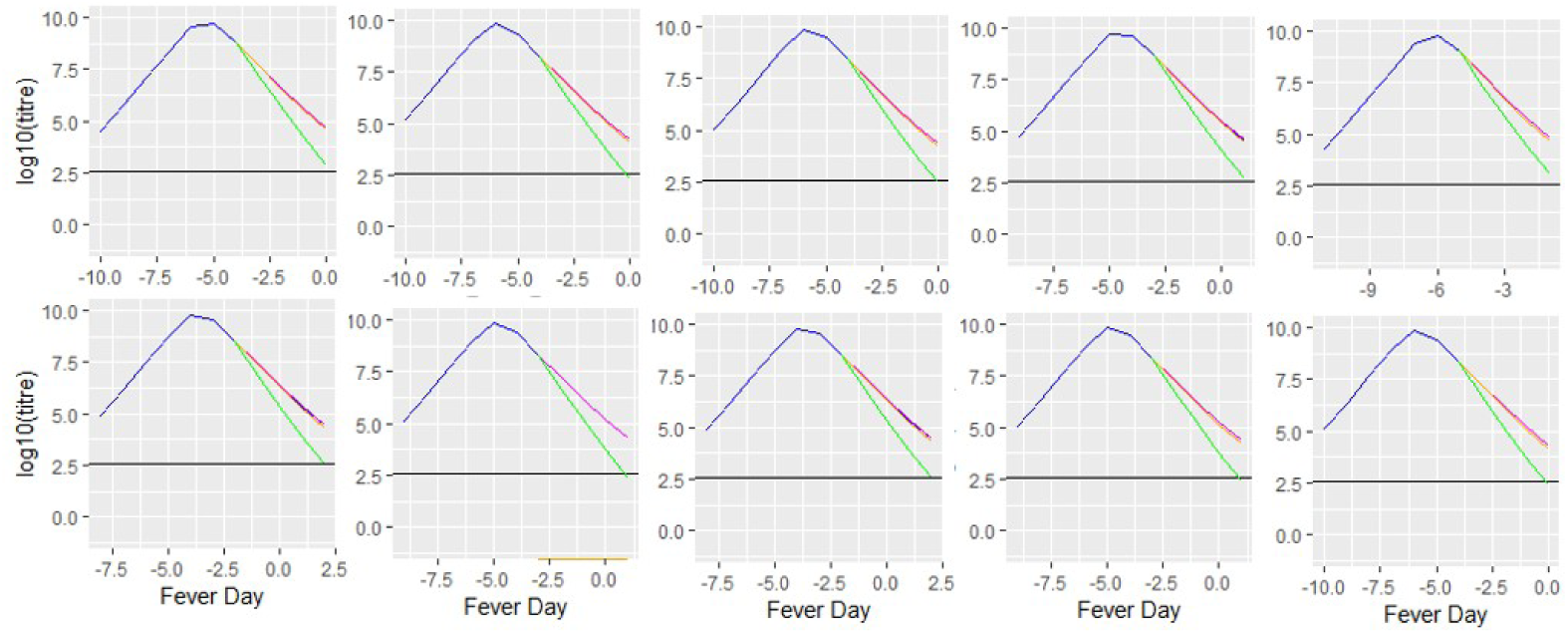
Effect of treatment of ten patients with DENV–TIPs doses on the day of reported symptoms when *σ* = 0.6. Subplots’ position and line colours have the same meaning as given in S6 Fig.

**S8 Fig.**
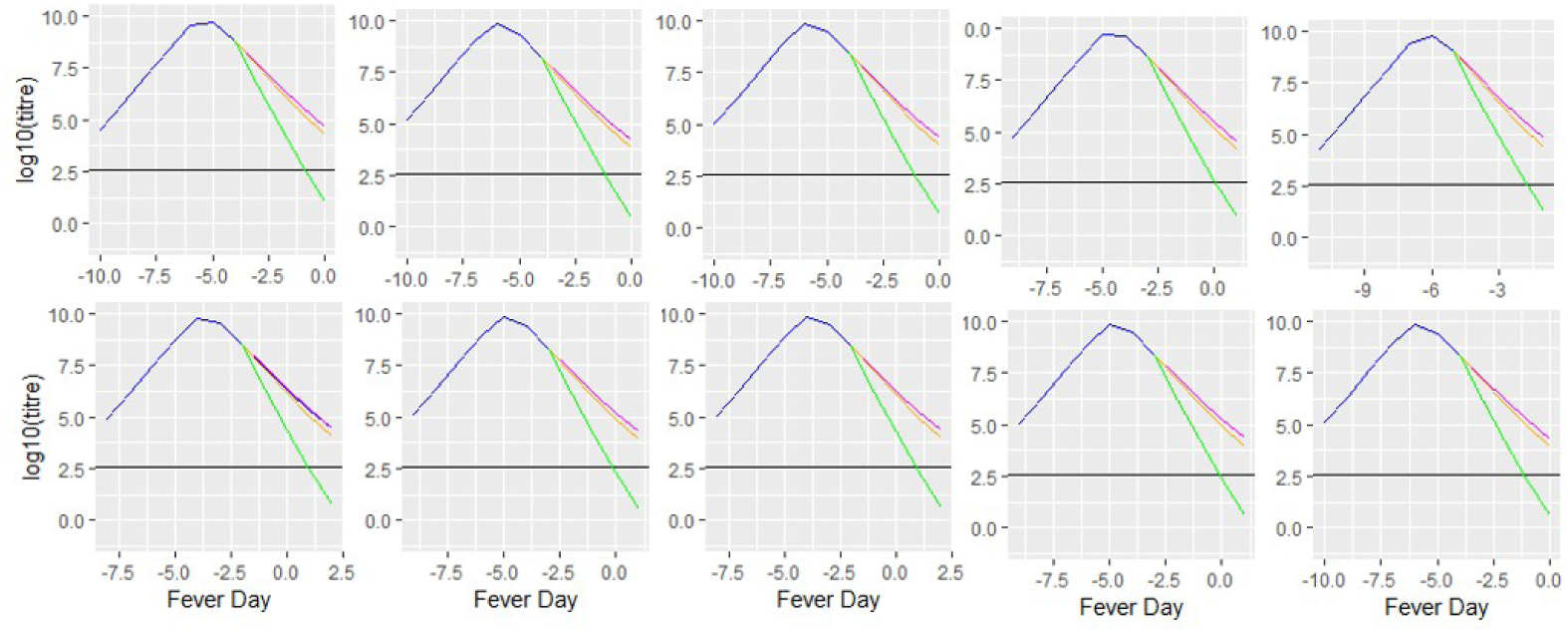
Effect of treatment of ten patients with DENV–TIPs doses on the day of reported symptoms when *σ* = 0.8. Subplots’ position and line colours have the same meaning as given in S6 Fig.

**S9 Fig.**
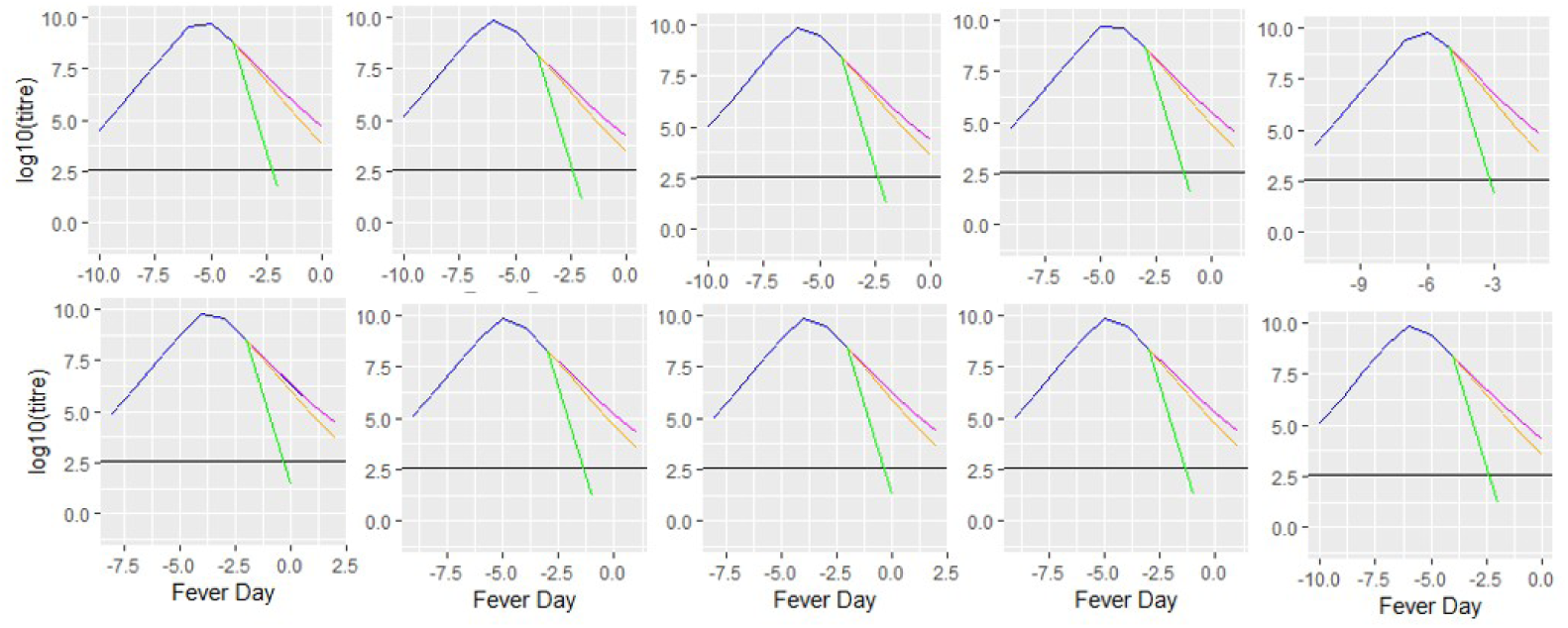
Effect of treatment of ten patients with DENV–TIPs doses on the day of reported symptoms when *σ* = 1, 0. Subplots’ position and line colours have the same meaning as given in S6 Fig.

**S10 Fig.**
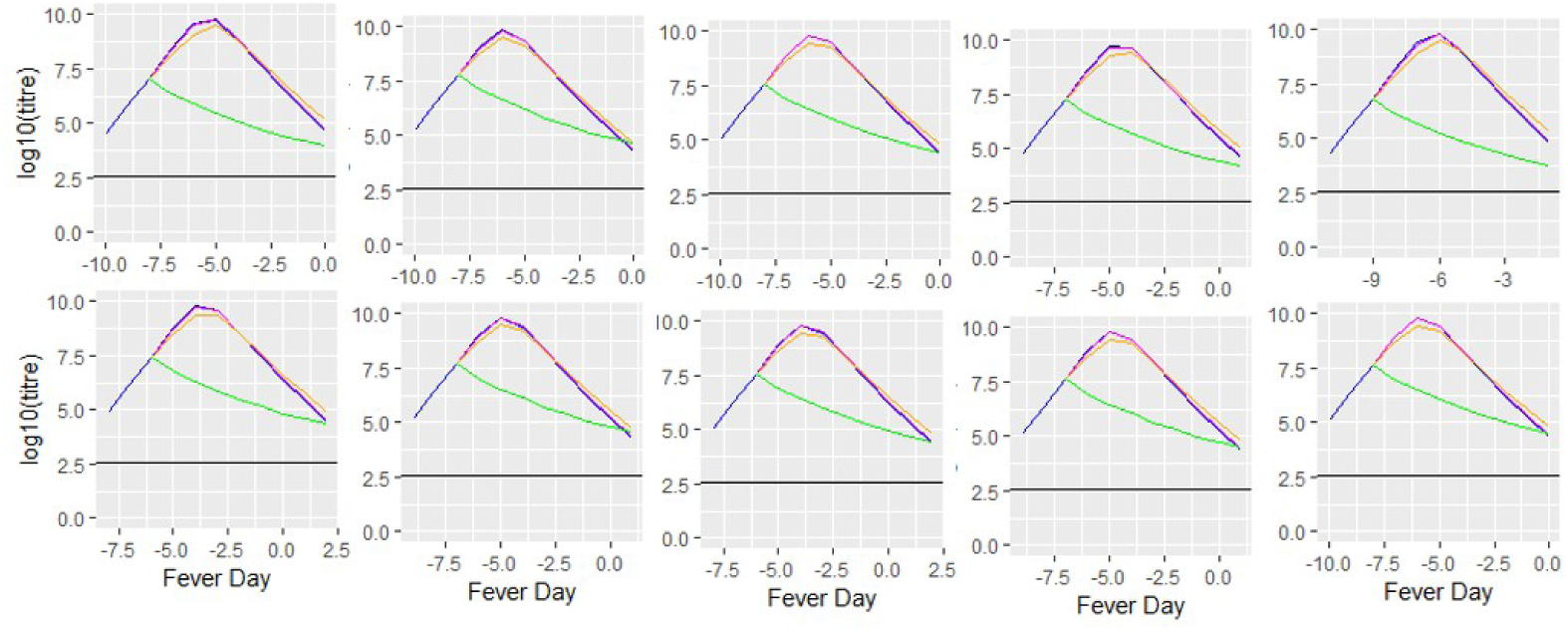
Effect of treatment of ten patients with DENV–TIPs doses on four days prior to the reported day of symptoms when *σ* = 0.7. Subplots’ position and line colours have the same meaning as given in S6 Fig.

**S11 Fig.**
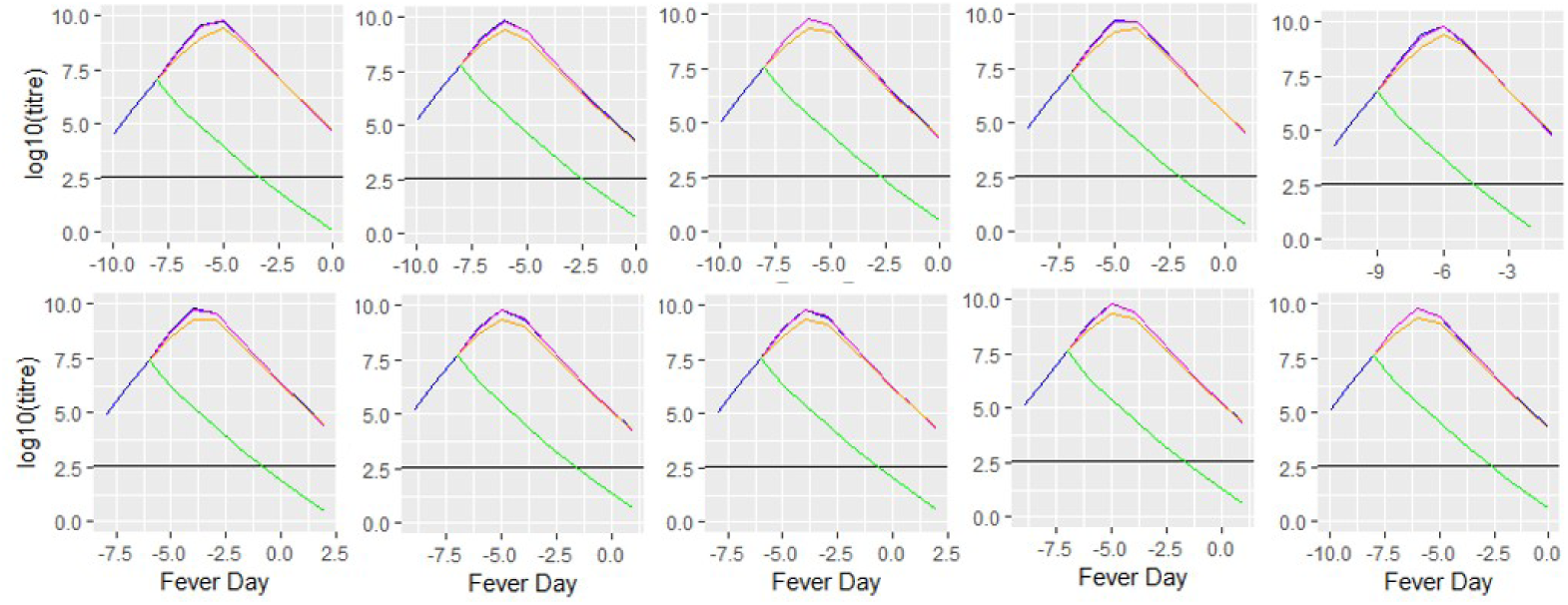
Effect of treatment of ten patients with DENV–TIPs doses on four days prior to the reported day of symptoms when *σ* = 0.8. Subplots’ position and line colours have the same meaning as given in S6 Fig.

**S12 Fig.**
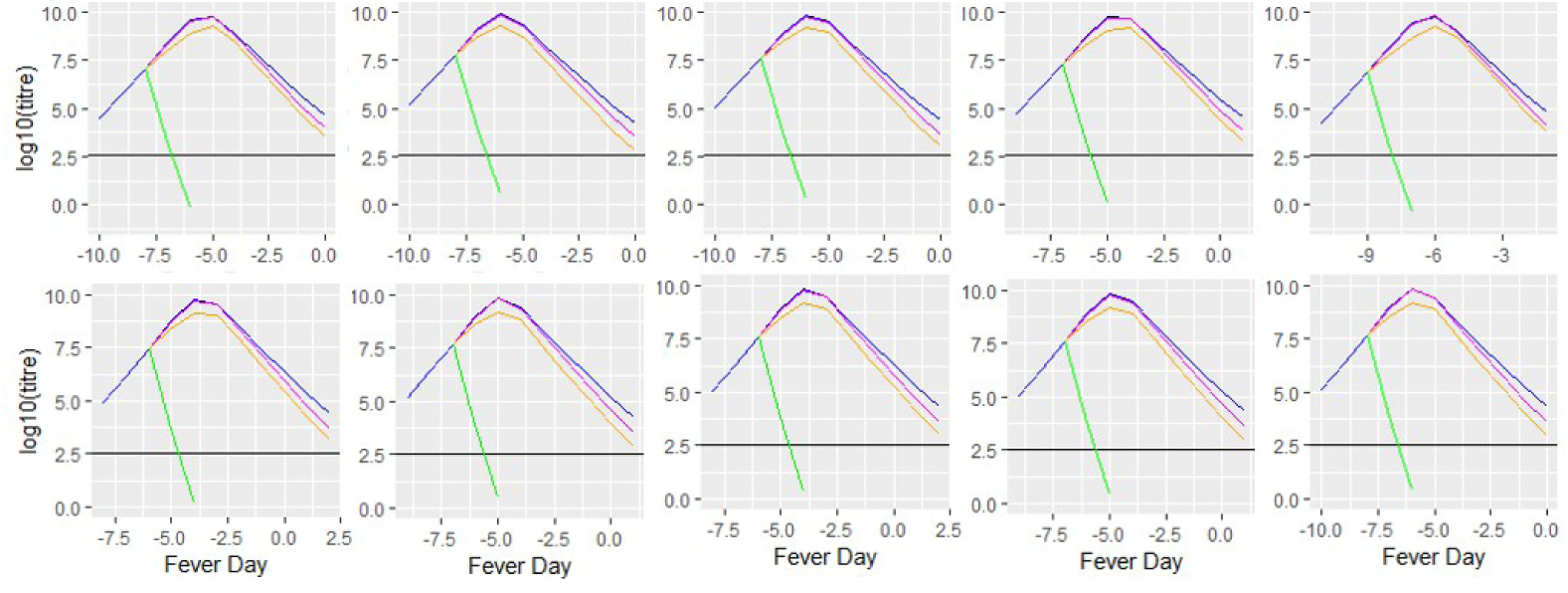
Effect of treatment of ten patients with DENV–TIPs doses on four days prior to the reported day of symptoms when *σ* = 1.0. Subplots’ position and line colours have the same meaning as given in S6 Fig.

**S13 Fig.**
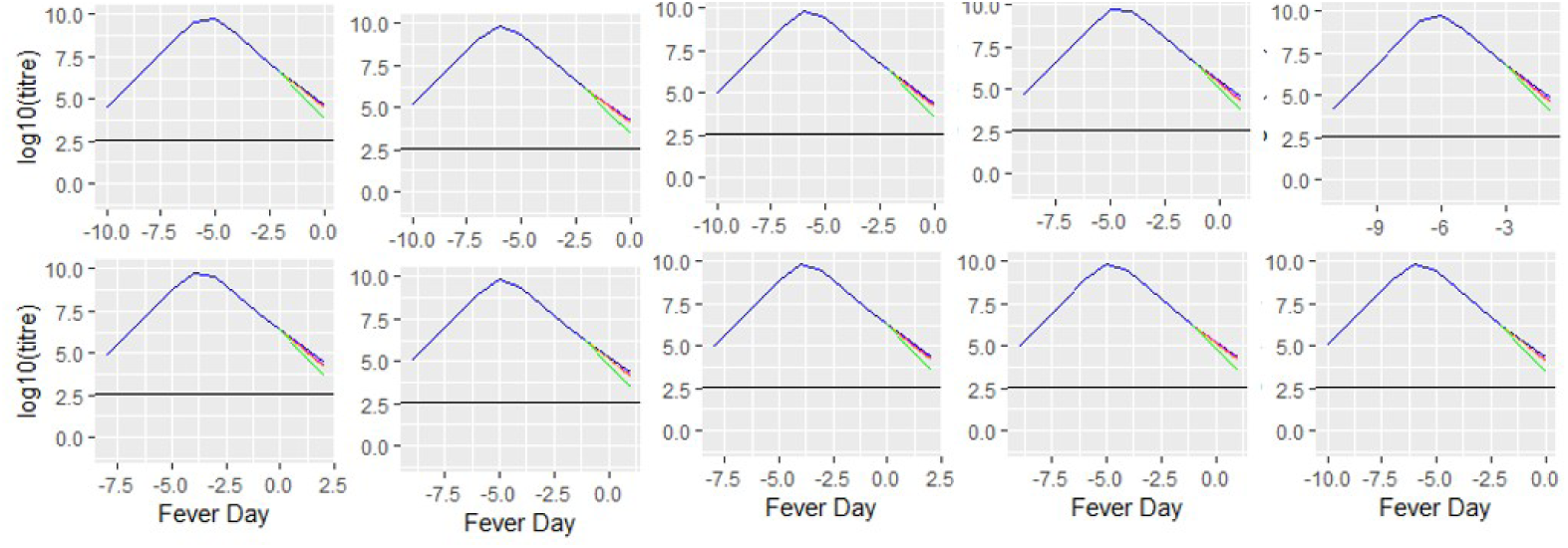
Effect of treatment of ten patients with DENV–TIPs doses on two days after the reported day of symptoms when *σ* = 0.5. Subplots’ position and line colours have the same meaning as given in S6 Fig.

**S14 Fig.**
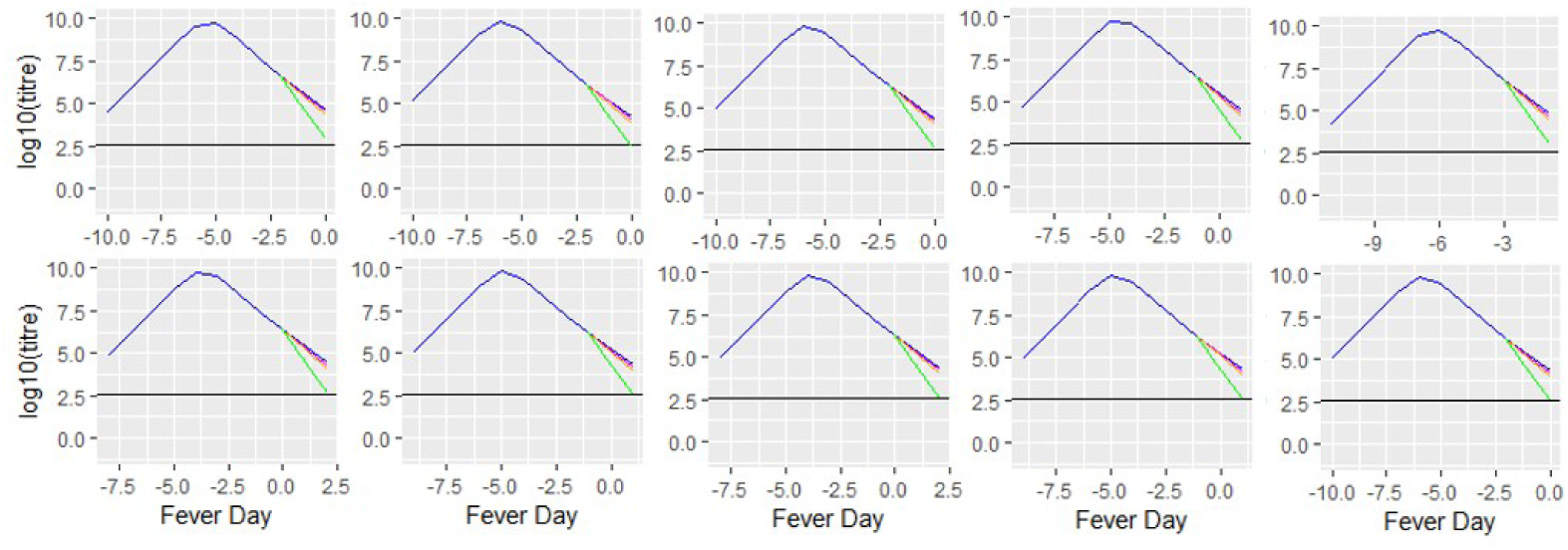
Effect of treatment of ten patients with DENV–TIPs doses on two days after the reported day of symptoms when *σ* = 0.7. Subplots’ position and line colours have the same meaning as given in S6 Fig.

**S15 Fig.**
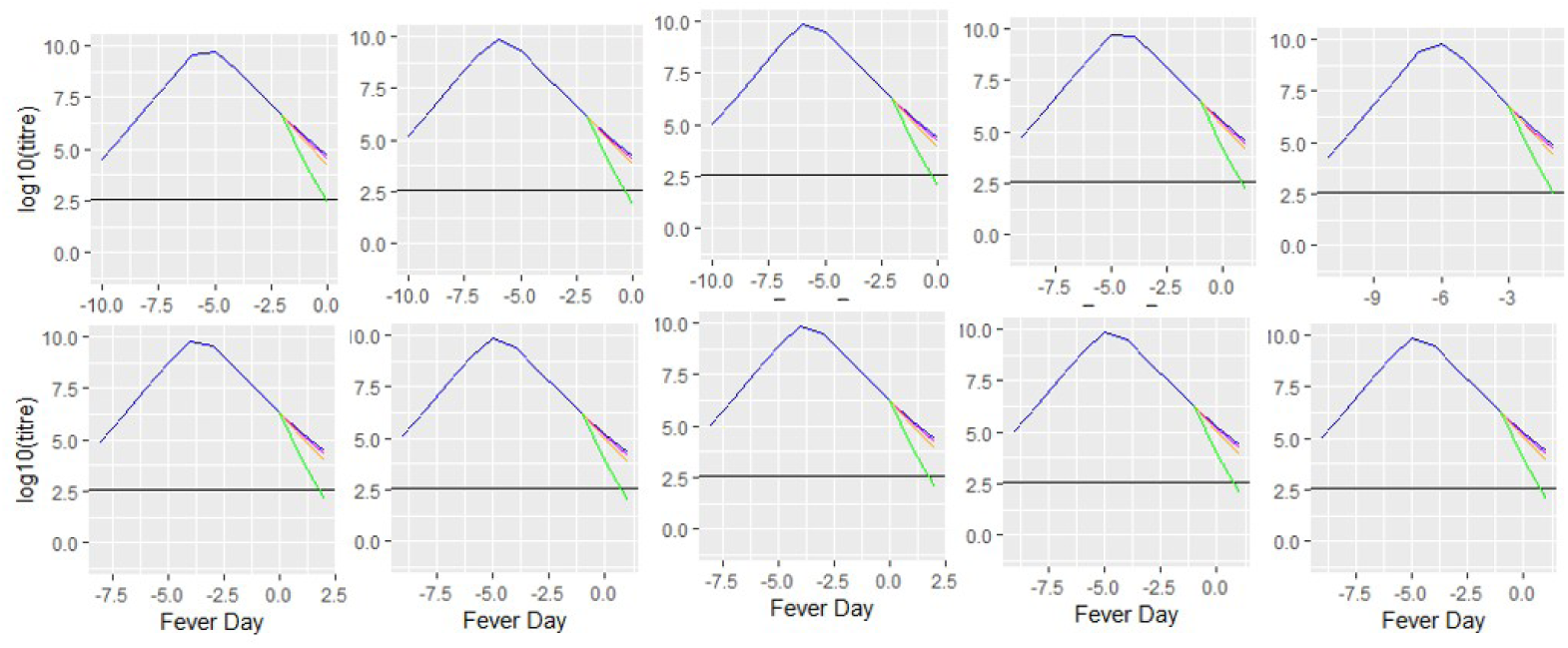
Effect of treatment of ten patients with DENV–TIPs doses on two days after the reported day of symptoms when *σ* = 0.8. Subplots’ position and line colours have the same meaning as given in S6 Fig.

**S16 Fig.**
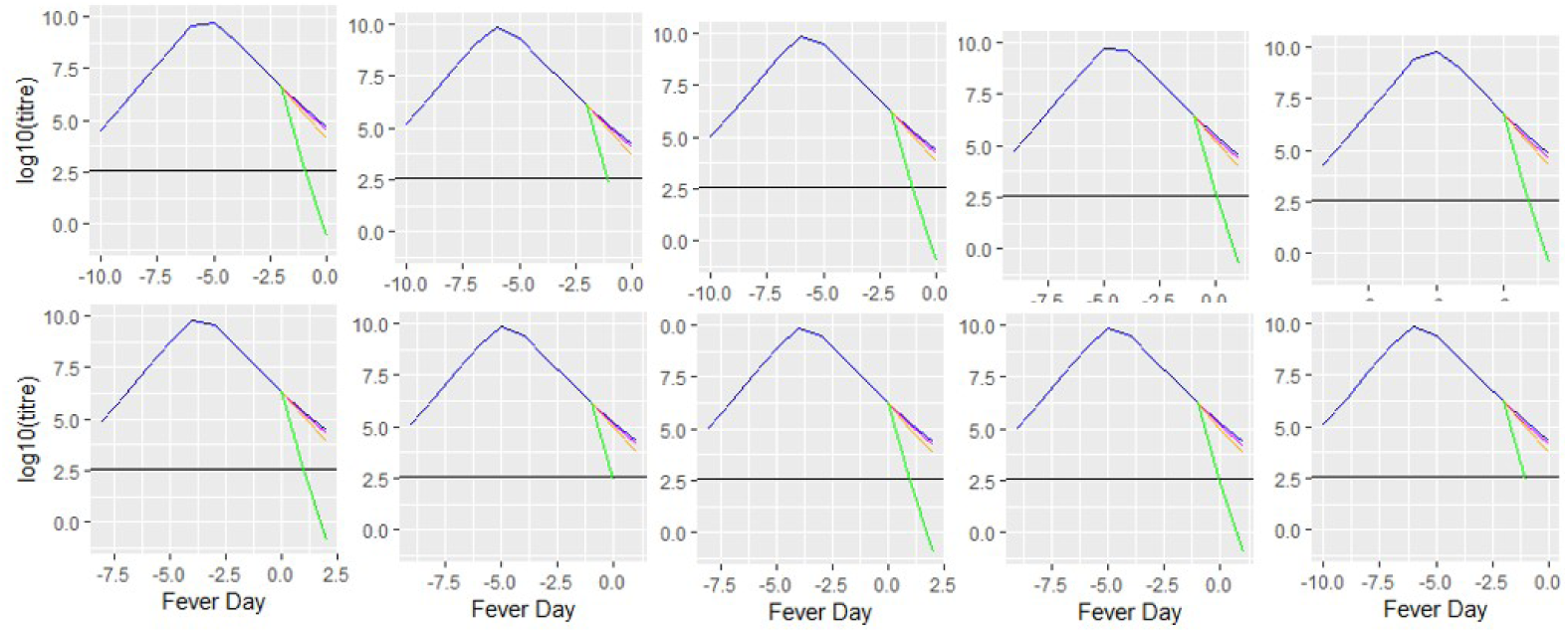
Effect of treatment of ten patients with DENV–TIPs doses on two days after the reported day of symptoms when *σ* = 1.0. Subplots’ position and line colours have the same meaning as given in S6 Fig.

**S1 File A compressed ZIP file consisting of: an R code file to run the Stan programs, plotting results and simulation studies; the Stan program for fitting model** *a* **to data; the Stan program for fitting model** *b* **to data; the data used to produce results in S5 Fig and S1 Table**.

## Acknowledgments

We thank Nguyet Minh Nguyen and co–authors for providing the data collected at the Hospital for Tropical Diseases, HCMC, Vietnam. We thank all those involved in the DARPA INTERCEPT Dengue TIPs project for their valuable suggestions and discussions. We thank the High Performance Computing (HPC) and Research Support of QUT for the computational resources.

